# All-trans Retinoic Acid Regulates miR-106a-5p Inhibition of Autophagic in Developing Cleft Palates

**DOI:** 10.1101/2020.11.05.369322

**Authors:** Lungang Shi, Yan Liang, Lijing Yang, Binchen Li, Binna Zhang, Congyuan Zhen, Jufeng Fan, Shijie Tang

## Abstract

**Background:** All-trans retinoic acid (atRA) results in cleft palate, but the cellular and molecular mechanisms underlying the teratogenic effects on palatal development have not been fully elucidated. Autophagy interruption has been reported to seriously affect embryonic-cell differentiation and development. This study aimed to verify whether atRA-induced cleft palate occurs because atRA blocks autophagy and stemness of embryonic palatal mesenchyme (MEPM) cells, which are maintained via the phosphatase and tensin homolog (PTEN)/protein kinase B (Akt)/mammalian target of rapamycin (mTOR) autophagic signaling pathway, and inhibits osteogenic-differentiation potential of MEPM cells, which could lead to the development of cleft palate.

**Methods:** To assess the stemness and pluripotency of MEPM cells, we analyzed their surfacemarkers using immunofluorescence (IF) and flow cytometry (FCM). Differentiation potentials, such as osteogenic, adipogenic, and chondrogenic differentiation, were induced. We also investigated the role of the PTEN/Akt/mTOR autophagic signaling pathway, which maintains the stemness and pluripotency of MEPM cells. Using transmission electron microscopy (TEM), Western blot analysis, quantitative reverse transcriptase polymerase chain reaction (RT-qPCR), messenger ribonucleic acid (mRNA) microarray, dual-luciferase reporter system, and exosomes, we found that atRA blocks autophagy and osteogenic differentiation of MEPM cells through micro-ribonucleic acid (miR)-106a-5p by targeting the PTEN/Akt/mTOR autophagic pathway.

**Results:** *In vitro* purified MEPM cells expressed cell surface markers similar to those of mouse bone marrow stem cells. Additionally, *in vitro* MEPM cells were ectomesenchymal and expressed the neural-crest marker human natural killer-1 (HNK-1), the mesodermal marker vimentin, and the ectodermal marker nestin. They were also positive for *in vitro* MEPM markers, including platelet-derived growth factor alpha (PDGFRα), ephrin B1 (Efnb1), odd-skipped related 2 (Osr2), and Meox2, as well as for stemness markers including POU class 5 homeobox 4 (Oct4), Nanog, and sex-determining region Y-related HMG box 2 (Sox2). MEPM cell pluripotency was retained through activation of the PTEN/Akt/mTOR autophagic signaling pathway. We found that atRA blocked MEPM cell pluripotency to inhibit osteogenic differentiation via miR-106a-5p targeting of PTEN mRNA and subsequent suppression of the PTEN/Akt/mTOR autophagic pathway.

**Conclusions:** *In vitro* cultured MEPM cells are ectomesenchymal stem cells that have strong osteogenic differentiation potential, and MEPM pluripotency is regulated by autophagy via the PTEN/AKT/mTOR signaling pathway. atRA disrupts MEPM cell pluripotency through PTEN/AKT/mTOR signaling inactivation where miR-106a-5p targets PTEN mRNA to reduce osteogenic differentiation of MEPM cells and results in the development of cleft palates. Our findings provide new insight into the mechanism underlying the development of cleft palate, and miR-106a-5p may act as a prenatal screening biomarker for cleft palate as well as a new diagnostic and therapeutic target.

## Introduction

Cleft palate is one of the most common congenital birth defects, and both genetic and environmental factors are involved^1^. The palate consists of the primary and secondary palates^2^, and cleft palate is caused by a failure in the formation of the secondary palate during embryonic development^3,4^. The development of the secondary palate is precisely regulated, including the growth, elevation, and fusion of the palatal shelves from embryonic day 13.5 (E13.5) to E15.5^5^. Failed elevation of the palatal processes is widely considered as one of the primary causes for failure in the formation of the secondary palate during embryonic development and ultimately results in cleft palate^6^. Palatal shelves are primarily composed of mesenchymal cells surrounded by an epithelium^3,7–9^. Thus, dysregulation of embryonic palatal mesenchyme (MEPM) cells, which includes reduced viability and differentiation potential, can cause cleft palate^10^.

All-trans retinoic acid (atRA), the oxidative metabolite of vitamin A, plays an important role in cell proliferation and differentiation as well as extracellular matrix production during embryonic development^11^. However, exogenous atRA is a potent teratogen during embryonic development to animals and humans^12,13^. In mouse embryos, cleft palate is a major malformation induced by atRA and is temporally dependent^13^. Accordingly, at E10, atRA exposure to pregnant mice induces small palatal shelves incapable of elevating, whereas atRA exposure at E12 results in normal-sized palatal shelves that contact but fail to fuse^14^. The molecular and cellular mechanisms for atRA regulation of MEPM cell proliferation and apoptosis have been widely studied, yet this does not fully explain the mechanism of cleft palate development. In addition to examining proliferation and apoptosis, more detailed characterization of atRA regulation of MEPM cells is required to understand the mechanisms of cleft palate development. Hence, the atRA mechanisms underlying cleft palate formation are complex and still not fully understood.

Autophagy plays an important role in embryonic development and differentiation^15,16^, critical to the elimination of misfolded protein aggregates, invading microorganisms, and damaged organelles^17,18^. Autophagy-defective organisms, including fungi, protozoa, worms, and insects, exhibit various abnormalities in differentiation and development^15^. Autophagic dysregulation can result in numerous human diseases including developmental disorders, neurodegenerative disease, and cancers^19^. However, the role of autophagy in cleft palate development is unknown.

For these reasons, we hypothesize that atRA impairs autophagy to cause cleft palate. Here, we characterize MEPM cells stemness and pluripotency and determine whether MEPM cells maintain pluripotency through the phosphatase and tensin homolog (PTEN)/protein kinase B (Akt)/mammalian target of rapamycin (mTOR) autophagic pathway. Further, we examine whether autophagy of MEPM cells is affected when exposed to atRA. We also determine how atRA exposure obstructs the PTEN/Akt/mTOR autophagic pathway by analyzing the expression of autophagic regulatory proteins including PTEN, Akt, and mTOR. We examine whether atRA exposure could regulate stemness and osteogenic differentiation of MEPM cells, and investigate autophagy and palatal shelf ossification in atRA-induced cleft palate from E13.5 to E17.5.To identify how atRA regulates the PTEN/Akt/mTOR autophagic pathway, we examine differentially expressed microRNAs in MEPM cells exposed to atRA by microarray, and we use bioinformatics methods to analyze differentially expressed microRNAs and their potential target genes. A dual-luciferase assay system is used to confirm the candidate microRNAs and their targets. We analyzed the targeted microRNA further to determine whether it was expressed in the exosomes secreted by atRA-treated-MEPM cells, which could act as markers for atRA-induced cleft palate.

## Materials and Methods

### Animals

All experiments involving animals were approved ethically by the Ethics Committee for Animal Experiments of Shantou University Medical College. one hundred Kunming mice were purchased from the Vital River Laboratory Animal Technology Co. Ltd. (Beijing, China). Mice were fed under a 12-house light/dark cycle, with standard food and tap water ad libitum. Females were crossed with males of similar weights and ages overnight, and on the following morning, a vaginal plug was considered as embryonic day 0.5 (E0.5).

### Primary cell culture

Primary mouse embryonic palatal mesenchymal (MEPM) cells were cultured as described elsewhere^20^. Cells from passages 0 – 2 were used.

### Clonogenic assays

The clonogenic assay was performed as a routine procedure. Briefly, in group1, we cultured 2×10^3^ MEPM cells per six-well plate in passage 1 (P1) in complete Dulbecco’s Modified Eagle’s Medium (DMEM) for 14daysin triplicate. In group 2, we treated 2×10^3^ MEPM cells per six-well plate in P1 with atRA(10 μmol/mL) for 24h, and then cultured the cells in complete DMEM for 14days in triplicate. The cell colonies that developed were fixed in 4% phosphate-buffered formalin and stained with a crystal violet solution (Solarbio Life Sciences, Beijing, China). Aggregates of ≥50 cells were counted as colonies.

### Immunofluorescence

MEPM cells were trypsinized at P0 and P1, resuspended, and inoculated in six-well plates at a density of 3×10^5^ cells/well in six-well plates (Corning, USA) for IF staining of MEPM cells using a standard protocol with antibodies (Table 1).

**Table 1.** Antibodies for Immunofluorescence and Western blot.

| Antibody | Manufacturer | Catalogue | Concentration |
| --- | --- | --- | --- |
| LC3A/B | Cell Signaling Technology | #12741 | 1: 100 |
| Vimentin | Abcam | ab92547 | 1: 100 |
| Nestin | Abcam | ab11306 | 1: 300 |
| HNK-1 | Abcam | sc81633 | 1: 100 |
| PDGFR-α | Abcam | ab203491 | 1: 500 |
| pan-keratin | CST | #4545T | 1: 300 |
| Efnb1 | Santa Cruz Bioscience | sc515264 | 1: 10 |
| Osr2 | Santa Cruz Bioscience | sc81971 | 1: 10 |
| Meox2 | Santa Cruz Bioscience | sc393516 | 1: 10 |
| RARα | Santa Cruz Bioscience | C-1 | 1:50 |
| LC3A/B | Cell Signaling Technology | #12741 | 1: 1000 |
| P62 | Cell Signaling Technology | #5114S | 1: 1000 |
| Nanog | Abcam | ab214549 | 1: 1000 |
| Oct4 | Abcam | ab200834 | 1: 1000 |
| Sox2 | Abcam | ab92494 | 1: 1000 |
| PTEN | Cell Signaling Technology | #9188T | 1: 1000 |
| Akt | Cell Signaling Technology | #9272 | 1: 1000 |
| mTOR | Cell Signaling Technology | #2972S | 1: 1000 |
| phospho-PTEN | Cell Signaling Technology | #9551T | 1: 1000 |
| phospho-Akt | Cell Signaling Technology | #4060T | 1: 1000 |
| phospho-mTOR | Cell Signaling Technology | #2971S | 1: 1000 |
| CD63 | ABclonal | A5271 | 1: 1000 |
| CD9 | Abcam | ab92726 | 1: 1000 |
| HSP70 | PROTEINTECH | 10995-1-AP | 1: 1000 |
| TSG101 | Abcam | Ab125011 | 1: 1000 |

### Histochemical staining

Embryonic heads were fixed, dehydrated and embedded by routine procedures. Histological sections (5 mm) were stained for immunohistochemistry using a standard protocol with antibodies for LC3A/B (1:200; #12741S; Cell Signaling Technology, USA).

### Goldner’s trichrome staining

Embryonic heads were fixed, dehydrated and embedded by routine procedures^21^. Histological sections (5 mm) were for Goldner’s trichrome staining using the manufacturer’s instructions to detect the ossification of the palate shelves with Goldner’s trichrome kits (Solarbio Life Sciences, Beijing, China).

### Flow cytometry

Cells were collected for flow cytometry (FCM) to identify the characteristics of MEPM cells using a standard protocol^22^. The mouse (MSC) analysis kit was obtained from Cyagen Biosciences, Inc. (Guangzhou, China). MEPM cells at passage 2 were harvested and re-suspended in 100 μL buffer solution (1% PBS supplemented with 0.1% FBS) at a cell density of 3×10^6^ cells/mL. Armenian hamster immunoglobulin G (IgG) isotype (2 μL) control antibody, purified anti-mouse Cluster of Differentiation 29 (CD29; 2 μL), rat IgG2b, κisotype control antibody (2 μL), anti-mouse CD44 (2 μL), anti-mouse CD117 (2 μL), rat IgG2a, κ isotype control antibody (2 μL), anti-mouse Sca-1 (2 μL), anti-mouse CD31 (2 μL), anti-mouse CD34 (2 μL), and anti-mouse CD90.2 (2 μL) were respectively added into and incubated for 30 min on ice. After two washes with buffer solution, the cells were re-suspended in 100 μL buffer solution. The cells were subsequently incubated with PE-conjugated goat anti-hamster lgG antibody (2 μL) and PE-conjugated goat anti-rat lgG antibody for 30 min on ice. After two washes with buffer solution, the cells were re-suspended in 400 μL buffer solution and the number of positive MEPM cells was quantified using a BD Accuri C6 flow cytometer (Becton-Dickinson, Franklin Lakes, NJ).

### Multi-lineage differentiation

Multi-lineage cellular differentiation was performed at P2 as previously described^20,23–25^, with the manufacturer’s instructions using osteogenesis, chondrogenesis, and adipogenesis assay kits (Cyagen Biosciences, Inc.).

### Adipogenic differentiation

Adipogenic induction was performed by culturing cells in six-well plates (Corning, USA), After cells reached confluence, they were cultured in mouse BMSC adipogenesis-inducing medium (AIM) for three days. Subsequently, the AIM was replaced with mouse BMSC adipogenesis-maintenance medium (AMM) for 24h, then switched back to AIM. After three cycles, the cells were cultured in the AMM for one week. Adipogenic differentiation was assessed with fresh Oil Red O solution staining as described elsewhere^26^.

### Chondrogenic differentiation

MEPM cells were grown in a monolayer. The cultured cells were maintained in mouse BMSC chondrogenic differentiation medium for four weeks with exchange every three days. Regular medium served as control. Immunohistochemical staining for collagen type II was stained to confirm the chondrogenic differentiation.

### Osteogenic differentiation

For osteogenic induction, MEPM cells were plated in six-well plates. After attainment of 80– 90% confluence, the cells were placed into mouse BMSC osteogenic-inducing medium for three weeks with the exchange every three days. Regular medium was used as a control. After a two-week-long induction, alkaline phosphatase (ALP) was stained using an ALP analysis kit (Solarbio LIFE SCIENCES) to confirm the osteogenic differentiation potency. ALP activity was performed as described elsewhere^27,28^. After a three-week-long induction, osteogenic induction was detected by Alizarin Red solution (ARS). ARS (Solarbio LIFE SCIENCES) (1%, pH4.2) staining and quantification were carried out as described elsewhere^29^.

### Quantitative real-time RT-PCR (RT-qPCR)

Total RNA was extracted using Trizol (Life Technologies) and then reverse transcribed into cDNA using a PrimeScript RT reagent kit (Takara). The PCR reaction system was prepared with a kit (SYBR Premix Ex Taq, Takara), and the reaction was performed on an Mx3000P real-time PCR machine (Agilent Stratagene, Santa Clara, CA, USA). The levels of target gene mRNA transcripts relative to control β-actin or gapdh were determined using the 2^−ΔΔCt^ method^30^. The RT-qPCR primer sequences are listed in Table 2.

**Table 2.**
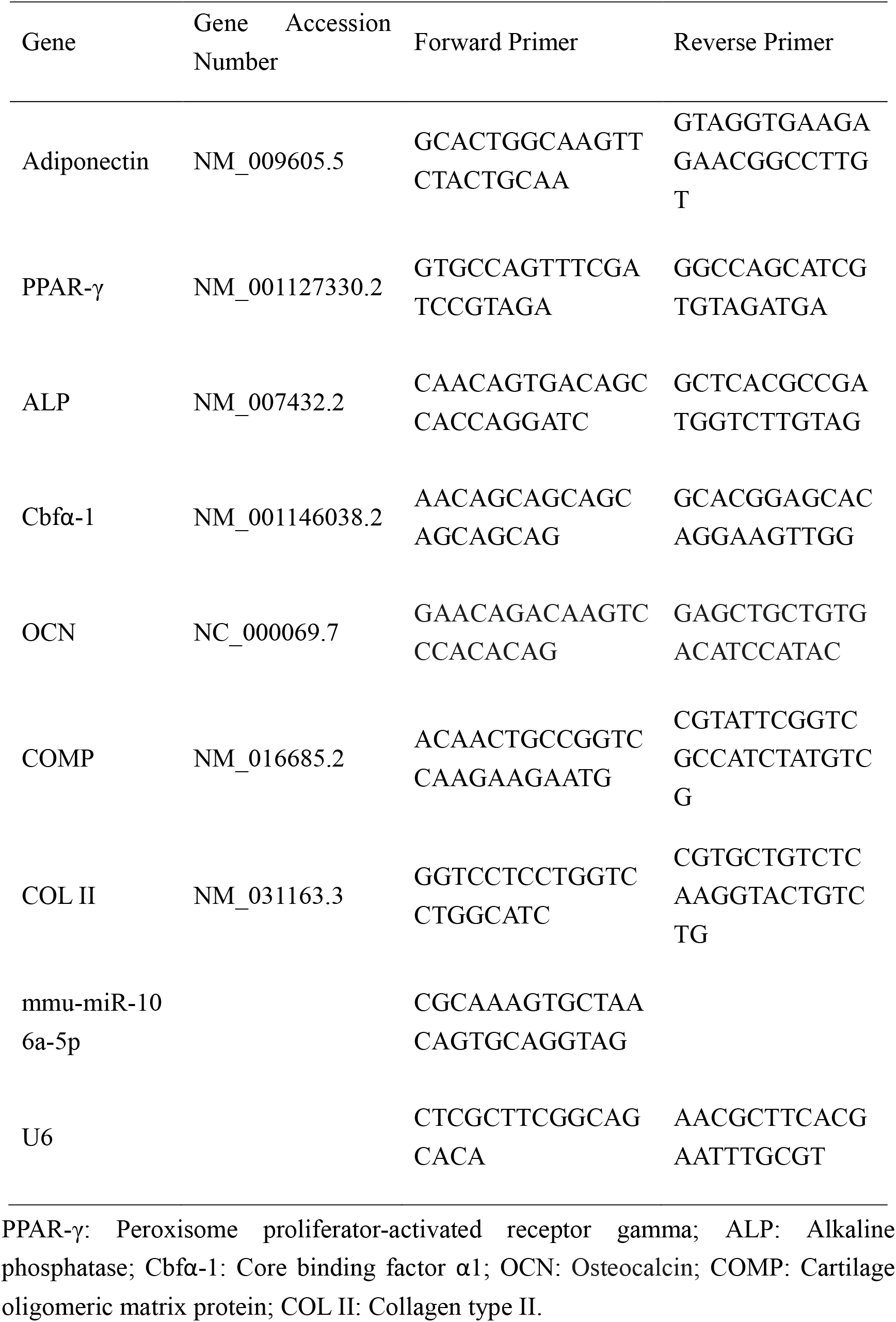
Primer sequences used for the relative quantification of the transcripts by RT-qPCR.

### Western blot analysis

The MEPM cells protein and palatal shelf protein extracts were analyzed by standard protocols^31^ with antibodies (Table1).

### Transmission electron microscopy (TEM)

Cells (5×10^4^–1×10^5^/condition) were centrifuged for 5 min at 4°C at 800 *g*. Subsequently, the supernatants were removed, the pellet was centrifuged at 4°C at 1200*g* for 10 min, and then fixed using Glutaraldehyde Fixed Solution for Electron Microscope (Pythonbio AAPR46, Pythonbiotech). We also fixed the palatal tissues using this same solution. Samples were then delivered to the Electron Microscopy Core Facility (ZHBY Biotech Co. Ltd, Nanchang China) for standard TEM analysis.

### Luciferase reporter assay

H293T cells were cultured in 24-well plates for 24h before transfection. The plasmid was constructed by Genechem (Shanghai, China). The Renilla reference plasmid, miR-106a-5p control cells, miR-106a-5p mimics cells, and the target gene 3’-UTR reporter gene plasmid were co-transfected. After transfection with Lipofectamine2000 (Invitrogen) for 48h, luciferase activity was examined using a double luciferase gene reporter kit (Genomeditech).

### MicroRNA microarray

RNA of sufficient quality from orofacial tissues of murine fetuses of E14.5 in the atRA group and the normal group (n = 3 per group) was submitted to KangChen-Biotech (Shanghai, China). The sample RNA was then marked with the miRCURY Array Power Labeling Kit (Exiqon, Denmark). The RNA was hybridized with the miRCURY LNA microRNA chip. The hybridization results were scanned by the Axon GenePix 4000B chip scanner, analyzed by GenePix pro V6.0. MEV software (v4.6, TIGR) was used for differentially expressed screening (with Fold Change ≥ 1.5, P-value <0.05) and cluster analysis.

### Isolation and identification of exosomes

Exosomes were isolated using ultracentrifugation^32^. In brief, MEPM cells culture supernatants were successively centrifuged at 800 g (10 minutes) and 2000 g (30 minutes) at 4°C. Then, exosomes were pelleted at 27300 rpm for 1 hour with an SW60 rotor (Beckman Coulter, California, USA). Exosome pellets were resuspended in phosphate-buffered saline (PBS) and centrifuged at 27300 rpm for 1 hour (SW60Ti rotor; Beckman Coulter). The exosome pellets were resuspended in PBS. Exosomes were identified using Nanosight 2000 analysis and transmission electron microscopy (TEM). Western blot analysis for exosomal markers, including CD9, CD63, Heat Shock Protein 70 (HSP70), and Tumor Susceptibility Gene 101 (TSG101), were used under standard protocols with antibodies (Table1). RNA from exosomes were extracted with a Total Exosome RNA Isolation Kit (Invitrogen, Carlsbad, CA, USA) for further analysis.

### Statistical analysis

The statistical analyses were performed using analysis of variance one-way ANOVA (SPSS 22.0) and Student *t*-tests. A p-value < 0.05 was considered statistically significant. All experiments were carried out in triplicate.

## Results

### Identification of fibroblastic MEPM cells from palatal shelves

The excised palatal shelves were cut into 1 mm^3^ cubes and plated in cell culture plates. Cells that migrated out of the palatal shelves were observed after 24h (Fig. 1A). The cells were collected by physical removal of palatal-shelf clumps and then digested, subcultured, and passaged every 3days. To characterize the *in vitro* migrating cells, we performed immunofluorescence (IF) staining for the mesodermal marker vimentin, the ectodermal marker nestin, the neural-crest marker HNK-1, and the epithelial-cell marker keratin. During the IF staining, the cells showed positive expression of vimentin and nestin; only 2% of cells stained positive for HNK-1, and 1% of cells stained positive for keratin (Fig. 1B). However, at passage 1 (P1), the cells that were positive for keratin at P0 lost keratin expression. In contrast, the cells that were positive for vimentin, nestin, and HNK-1 remained positive (Fig. 1B). These *in vitro* cultured cells also stained positive for MEPM cell markers including platelet-derived growth factor alpha (PDGFRα), ephrin B1 (Efnb1), odd-skipped related 2 (Osr2), and Meox2 (Fig. 1C). These results demonstrated that the cells isolated from palatal shelves were indeed MEPM cells, and cell passaging enables the purification of MEPM cells *in vitro*.

**Fig. 1.**
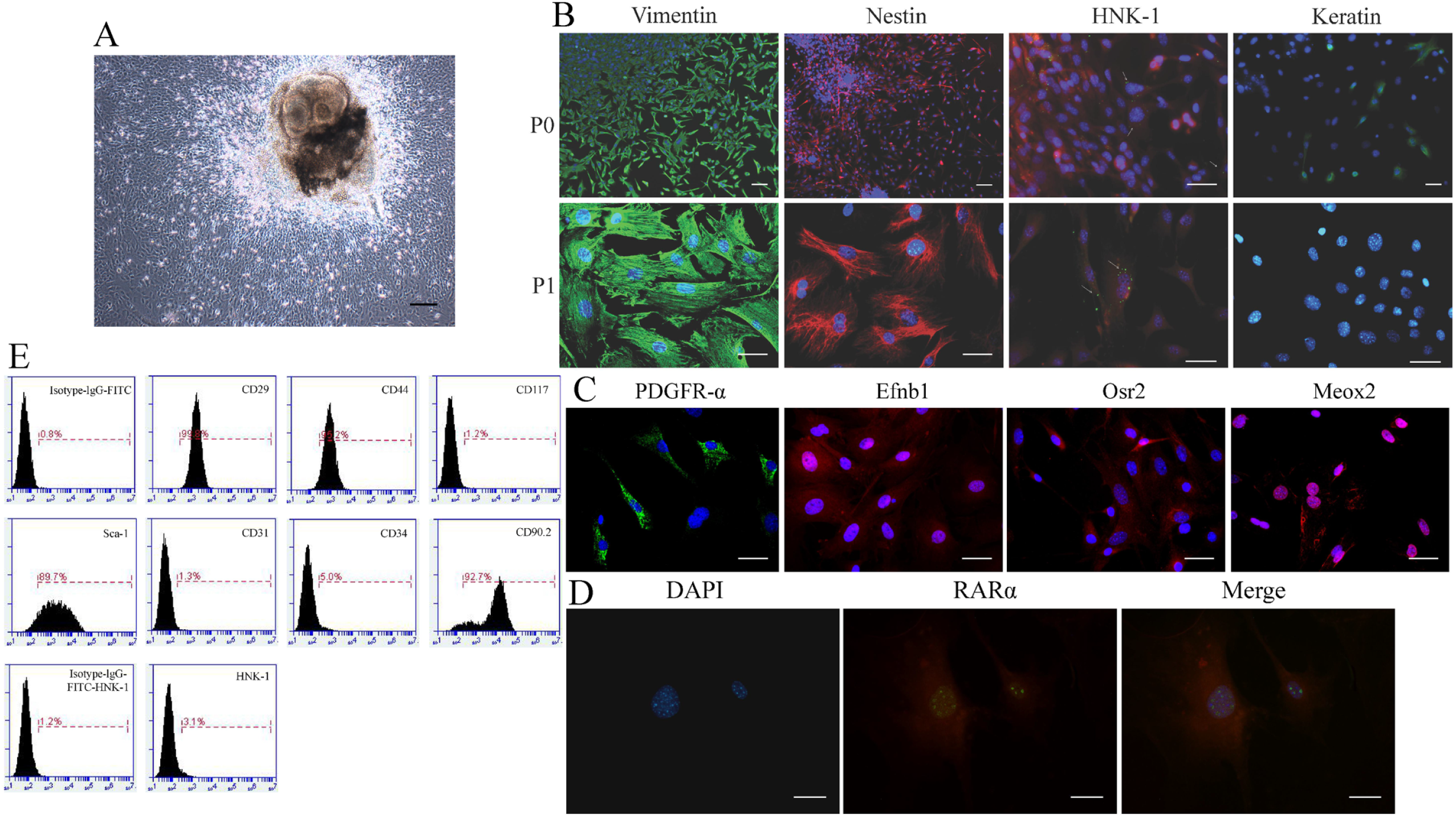
Characterization of primary mouse embryonic palatal mesenchymal (MEPM) cells. (A) Morphology of MEPM cells emerging from palatal shelves. MEPM cells appear as fibroblasts morphologically. Scale bar: 100 μm. (B) Immunofluorescence (IF) staining for vimentin (green), nestin (red), HNK-1 (green), and keratin (green). (C) IF staining for PDGFRα (green), Efnb1 (red), Osr2 (red), and Meox2 (red). (D) IF staining for RARα (green). (D) MEPM cell surface markers were characterized using flow cytometry. Cell nuclei (blue) were counter stained with 4’,6’-diamidino-2-phenylindole (DAPI) and the cytoplasm (red) was Evan blue stained with vimentin and nestin. Scale bar: 20 μm.

Colony formation capacity was an important indicator of stemness. We found large numbers of colony-forming units after 14 days of culture of MEPM cells (Fig. 4F). Expression of Oct4, Nanog, and Sox2 is essential for maintaining stem cell pluripotency^33–35^. IF staining confirmed that MEPM cells were positive for all three of these markers (Fig. 4E). Furthermore, FCM results demonstrated that MEPM cells had high expression levels of CD29, CD44, CD90.2, and stromal cell antigen (Stro-1), while CD34 exhibited low expression. However, the *in vitro* cultured cells did not express BMSC markers CD31 and CD117 (Fig. 1E). Only 1.9% of the cells stained positive for HNK-1, consistent with the results reported by Gazarian *et al*^36^.

Therefore, the expression profiles for cell surface and stemness markers, as well as the clonogenic assays, demonstrated that *in vitro* purified MEPM cells were ectomesenchymal stem cells by nature.

### Multi-lineage differentiation potential of MEPM cells

To examine the pluripotency of the *in vitro* cultured MEPM cells, we performed differentiation protocols for adipogenic, osteogenic, and chondrogenic cell lineages.

After culture in adipogenic medium for 3 weeks, approximately 20% of MEPM cells stained positive for Oil Red O. In contrast, about 1% of cells in the control group stained positive for Oil Red O (Fig. 2A). Adipogenic differentiation was further verified by RT-qPCR analysis. Adipogenic-related genes, Leptin and PPAR-γ, expression in cells treated with the adipogenic medium were significantly higher than those in the control group (Fig. 2A), demonstrating that MEPM cells had the potential of adipogenic differentiation.

**Fig. 2.**
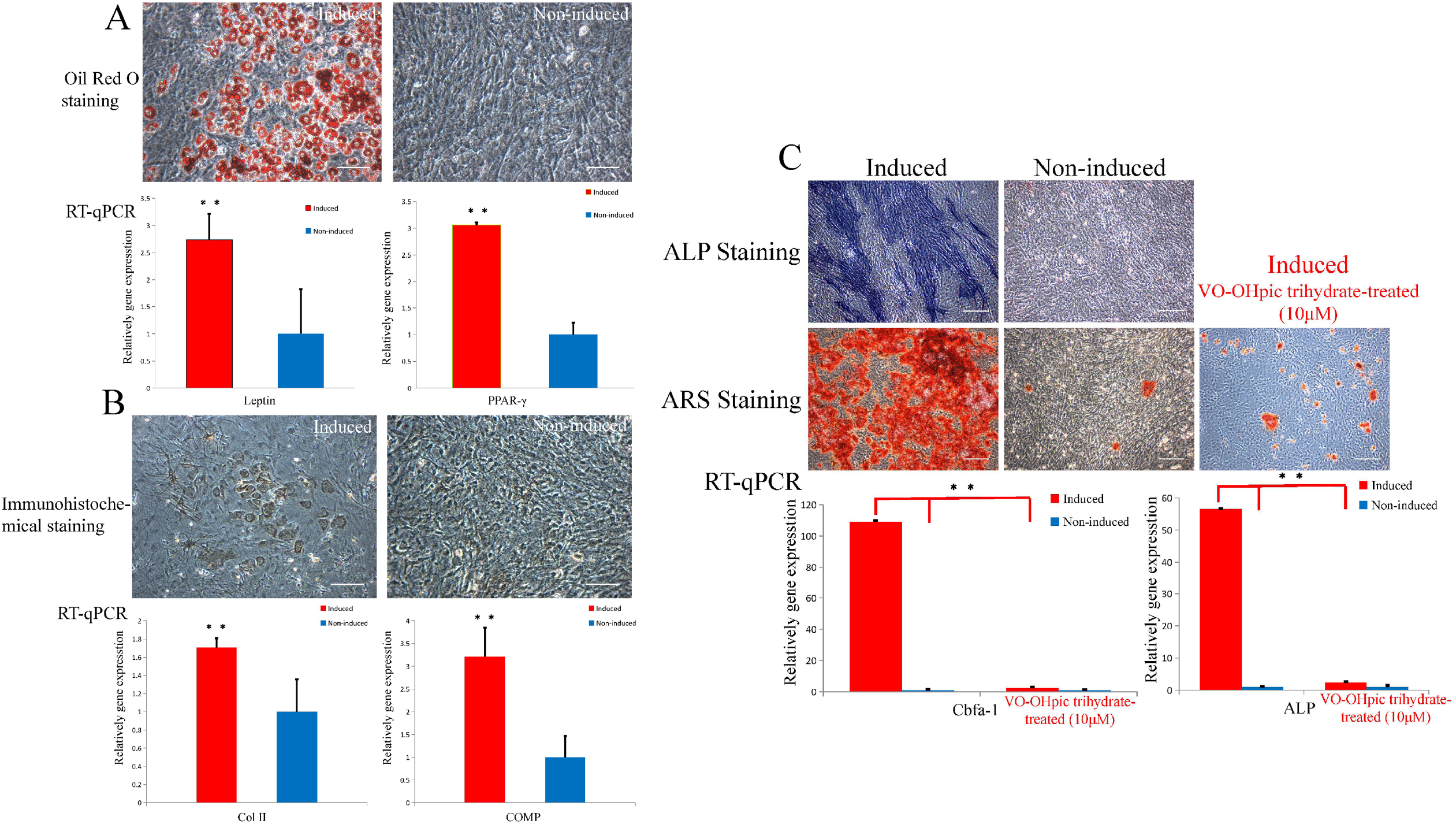
Multi-lineage differentiation of MEPM cells. (A) Adipogenic differentiation of MEPM cells, adipogenic-inducing groups (left panel) were positive for Oil Red O staining and negative in non-induction groups (right panel). Results of RT-qPCR showed the upregulation of Leptin and PPAR-γ in the induced compare to non-induction groups. Mean (± standard deviation) values from three independent experiments are shown. (B) Osteogenic differentiation of MEPM cells. After 2 weeks of osteogenic induction, the induced group (left panel) showed positive staining for alkaline phosphatase (ALP), and after 4 weeks of osteogenic induction, Alizarin Red staining (ARS) showed intensive calcium deposition in the induced group (left panel) as against very weak staining in the non-induced group (right panel), Results of RT-qPCR showed upregulation of ALP and Cbfα-1 in the induced group relative to non-induced groups. Mean (±standard deviation) values from three independent experiments are shown. (C) Chondrogenic differentiation of MEPM cells, immunohistochemical staining showing positive and negative staining for collagen type II (Col II) in the induced (left panel) compare to non-induced groups (right panel), respectively, results of RT-qPCR showing upregulation of COMP and Col II in the induced group compared to non-induced groups. Mean (±standard deviation) values from three independent experiments are shown. Scale bar: 50 μm; N = 3; *P < 0.05; **P < 0.01.

The MEPM cell chondrogenic differentiation potential was detected by immunohistochemical staining for collagen type II. Approximately 5% of cells in induction groups showed positive expression of collagen type II (Fig. 2B).Chondrogenic differentiation potential was further confirmed by RT-qPCR analysis. Collagen type II and the cartilage oligomeric matrix protein (COMP) were highly expressed in MEPM cells subjected to chondrogenic induction (Fig. 2B). Hence, these results indicated that *in vitro* purified MEPM cells have chondrogenic differentiation potential.

After 2 weeks of osteogenic induction, over 25% of cells stained positive for ALP compared to 2%of cells in the control group (Fig. 2C). The induced osteogenic phenotype was further confirmed by Alizarin Red staining, demonstrating intracellular calcium deposition. Approximately 93% of cells showed mineralized nodules after 4 weeks of osteogenic induction (Fig. 2C). The large extent of mineralized nodules indicated that *in vitro* purified MEPM cells possessed high osteogenic differentiation potential. The expression of osteogenic-related genes was verified using RT-qPCR, ALP and core binding factor α1 (Cbfα-1) mRNA levels were upregulated (Fig. 2C). These results indicated the strong osteogenic potential of *in vitro* MEPM cells.

### Characterization of autophagy in MEPM cells

The role of autophagy in stem cell survival and pluripotency has been extensively studied in recent years^37^. Studies show that autophagosome levels are higher in MSCs than in many differentiated cell types^38^. However, it is not clear whether undifferentiated MEPM cells also exhibit a high level of autophagy. In this study, to investigate the capacity of autophagy in MEPM cells, we cultured MEPM cells under differentiation conditions and assayed levels of LC3, type II (LC3-II), a biochemical marker for autophagy^39^. IF staining for LC3-II revealed the presence of numerous autophagosomes in MEPM cells prior to differentiation (Fig. 3A). However, the presence of autophagosomes was significantly reduced following osteogenic differentiation (Fig. 3A).

**Fig. 3.**
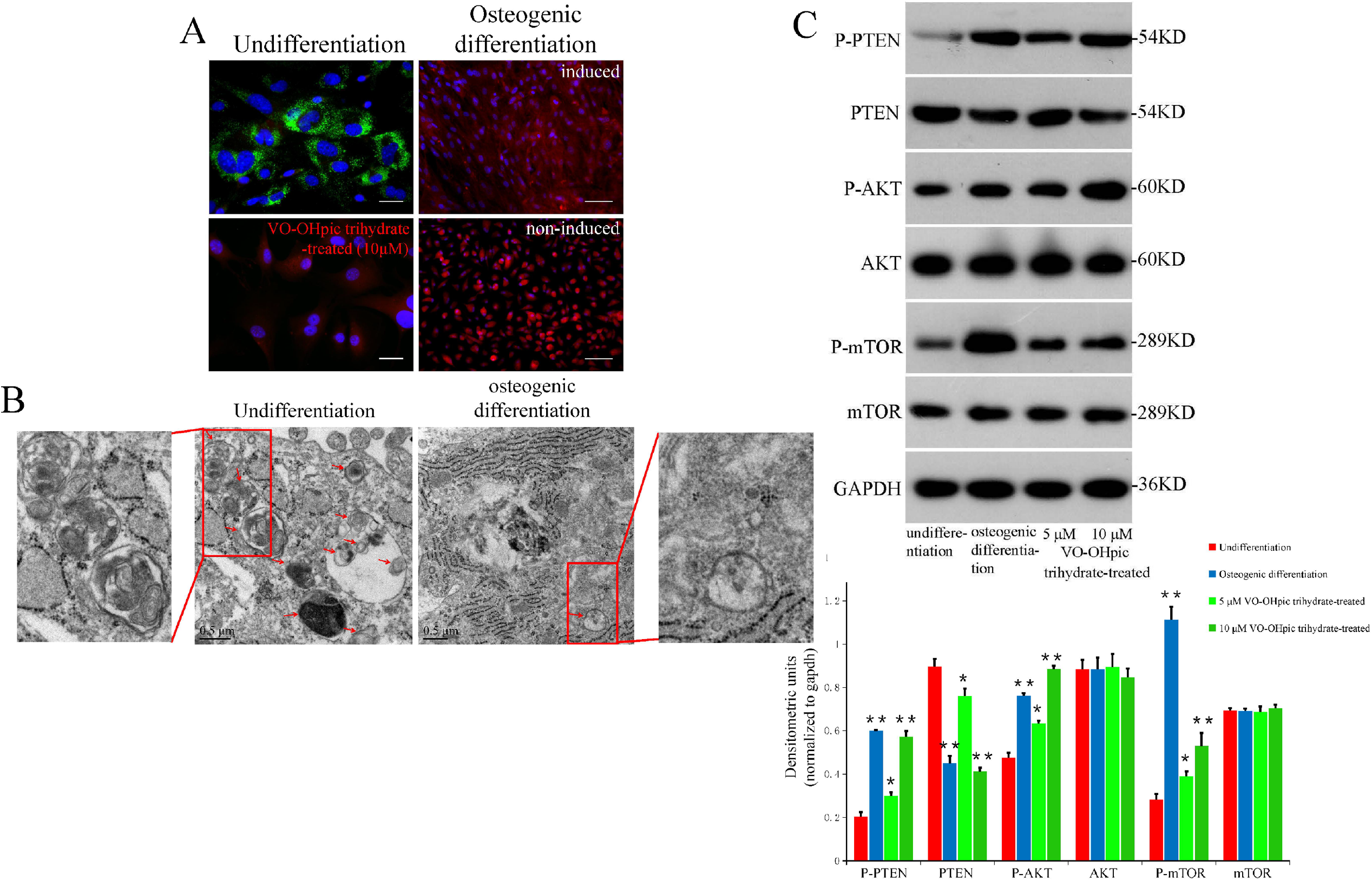
Characteristics of autophagy in MEPM cells. (A) IF staining showed positive staining for LC3-II before differentiation and negative staining after osteogenic, adipogenic differentiation, control groups after 3-weeks of culture, at passage 4 and MEPM cells exposed to VO-OHpic trihydrate. Cell nuclei (blue) were counterstained with 4’,6’-diamidino-2-phenylindole (DAPI). Cytoplasm(red) was stained with Evan blue. Scale bar: 20 μm. (B) Transmission electron microscopy image showing large numbers of autophagosomes (red arrows) before induction, which decreased sharply after osteogenic differentiation. (C) After 4 weeks of osteogenic induction MEPM cells at passage 2 exposed to VO-OHpic trihydrate, Alizarin Red staining (ARS) showed weak calcium deposition in the VO-OHpic trihydrate-treated group (left panel), and negative staining in the non-induced group (right panel). Scale bar: 50μm. Results of RT-qPCR showing upregulation of ALP and Cbfα-1 in the induced group compared to the non-induced and VO-OHpic trihydrate treated groups. (**P < 0.01). (D) Western blot analysis of phosphorylated and non-phosphorylated PTEN, AKT, and mTOR. N = 3; *P < 0.05; **P < 0.01.

These results were further verified by TEM, in which undifferentiated MEPM cells showed large numbers of autophagosomes; upon osteoblast differentiation of MEPM cells *in vitro*, these numbers were reduced (Fig. 3B). These results showed that high basal levels of autophagy were characteristic of undifferentiated MEPM cells.

Therefore, these results confirmed the role of autophagy in maintaining MEPM cell pluripotency.

### PTEN/Akt/mTOR autophagy signaling regulates MEPM cell pluripotency

The PI3K/Akt/mTOR pathway plays an important role as a regulator of autophagy^40^. PTEN, a tumor suppressor that suppresses activation of the PI3K/Akt/mTOR signaling pathway, has been suggested to activate autophagy^41^. To explore whether MEPM cells maintain pluripotency through the PTEN/Akt/mTOR autophagy signaling pathway, we examined pan- and phosphorylated-PTEN (phospho-PTEN), Akt, and mTOR. Western blot showed that the expression level of phospho-PTEN increased after osteogenic differentiation, whereas total PTEN protein expression was sharply downregulated. Consistent with the altered PTEN level, the downstream molecules for the phosphorylation of AKT and mTOR-two pivotal inhibitors of autophagy were upregulated without a concomitant effect on Akt and mTOR total protein expression (Fig. 3C). In contrast, prior to differentiation, phospho-PTEN levels were low, and total PTEN was highly expressed; however, phospho-AKT and phospho-mTOR exhibited low expression levels (Fig. 3C). Furthermore, to validate the our conclusion and ensure the specificity of these pathways, MEPM cells were treated with VO-OHpic trihydrate, the antagonist of PTEN, at a dose of 10 μM for 24h. The PTEN-Akt-mTOR signaling protein was then analyzed via Western blot. Total PTEN protein expression was restrained and downregulated dramatically, those of phosphorylated-PTEN, -Akt, and -mTOR levels were upregulated considerably, and total expression of AKT and mTOR remained unchanged (Fig. 3C). Similarly, IF staining for LC3-II demonstrated that VO-OHpic trihydrate exposure (10 μM) to *in vitro* MEPM cells for 24h resulted in autophagosome reduction (Fig. 3A). Furthermore, we examined the osteogenic-differentiation potency of MEPM cells that were pretreated with 10 μM VO-OHpic trihydrate for 24h. After 4 weeks of induction, Alizarin Red staining, RT-qPCR of ALP and Cbfα-1 results showed that VO-OHpic trihydrate treatment of MEPM cells *in vitro* could reduce their osteogenic differentiation potential considerably (Fig. 2C). In conclusion, these results demonstrated that MEPM cells maintained pluripotency through the PTEN/Akt/mTOR autophagic signaling pathway.

### atRA significantly reduces autophagosome levels of MEPM cells

Cultured MEPM cells at P1 were treated for 24 h with the following doses of atRA: 3, 5, 7, 10, and 15 μmol/mL. IF staining for LC3-II showed that autophagosome levels decreased as the dose of atRA increased (Fig. 4A). At a dose of 10 μmol/mL atRA, the autophagosomes disappeared. These findings were verified by Western blot; the protein ratio of LC3-II to LC3-I was greatly decreased. Moreover, the levels of the autophagy adaptor protein p62, which is degraded by autophagy and autophagic flux, were increased (Fig. 4B). Furthermore, TEM showed a largenumber of autophagosomes in undifferentiated MEPM cells, but treatment with 10 μmol/mL atRA reduced the number of autophagosomes (Fig. 4C). Thus, these results showed that atRA affects autophagy of MEPM cells *in vitro*.

**Fig. 4.**
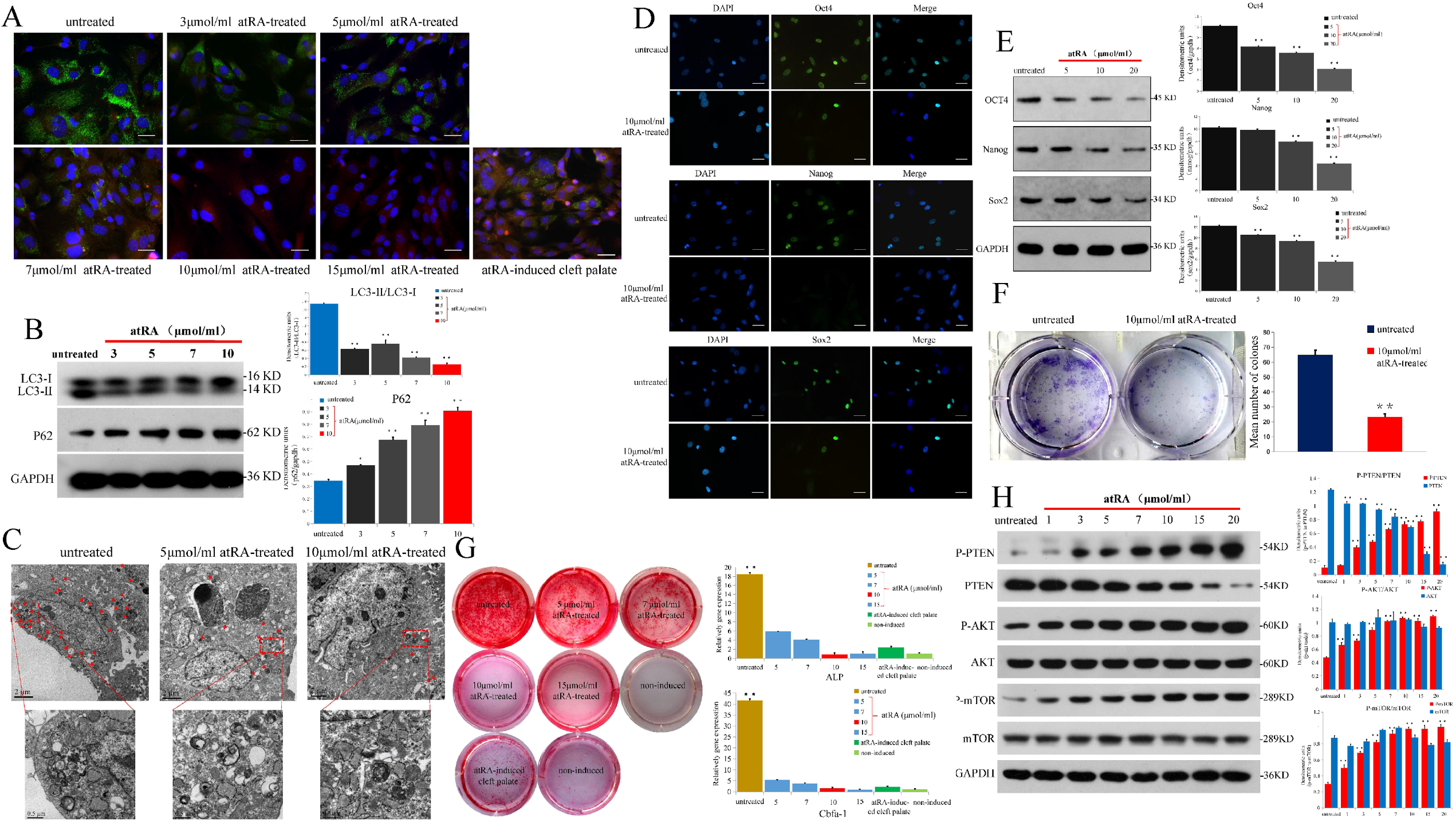
Characteristics of MEPM cells treated with atRA. (A) IF staining for LC3-II is weakly positive when treated with different doses of atRA and MEPM cells separated from atRA-induced cleft palate. Scale bar: 20 μm. (B) Results of Western blot analysis showing expressions of LC3-II/I and p62 proteins before and after atRA-treated. (C) Transmission electron microscopy image showing decreased sharply of autophagosomes (red arrows) after atRA-treated. (D) IF staining for Oct4 (green), Nanog (green), and Sox2 (green). Scale bar: 20 μm. (E) Western blot analysis of protein levels of Oct4, Nanog, and Sox2. (F) Clonogenic assays of MEPM cells treated with atRA (10 μmol/mL). (G) After 4 weeks of osteogenic induction, Alizarin Red staining (ARS) showed weak calcium deposition in the induced group (left panel), negative staining in the non-induced group (right panel) and MEPM cells separated from atRA-induced cleft palate. Scale bar: 50 μm. (H) Results of RT-qPCR showing upregulation of ALP and Cbfα-1 in the induced group compared to non-induced groups and in MEPM cells separated from atRA-induced cleft palate. (I) atRA blocked the PTEN/Akt/mTOR autophagy signaling pathway, according to Western blot analysis of protein levels of phosphorylated and nonphosphorylated PTEN, Akt, and mTOR. N = 3; *P < 0.05; **P < 0.01.

### Autophagosome reduction in atRA-treated MEPM cells inhibits stemness and osteogenic differentiation

When undifferentiated MEPM cells were treated with 10 μmol/mL atRA for 24h, the number of autophagosomes was dramatically reduced. When we explored changes in stemness of undifferentiated MEPM cells treated with 10 μmol/mL atRA for 24h, we found that numbers of colony forming units were greatly reduced after 14 days of culture (Fig. 4F). MEPM cell IF results also showed that expression of stemness markers, including Oct4, Nanog, and Sox2, were reduced dramatically (Fig. 4D). Western blot results showed that Oct4, Nanog, and Sox2 proteins were downregulated when undifferentiated MEPM cells were treated with atRA (5,10, and 20μmol/mL) for 24h, compared with untreated undifferentiated MEPM cells (Fig. 4E). Therefore, we concluded that atRA could impair the stemness of MEPM cells.

To further confirm this finding, we explored the osteogenic-differentiation potency of MEPM cells treated with the following doses of atRA for 24h: 5, 7, 10, and 15 μmol/mL. After 4 weeks of osteogenic induction, Alizarin Red staining showed a reduction in mineralized nodules in MEPM cells with 5, 7, 10, and 15 μmol/mL of atRA (Fig. 4G). This was further verified by RT-qPCR analysis; osteogenic-related genes, ALP and Cbfα-1, showed a significant decrease in mRNA levels in the atRA-treated MEPM cells (Fig. 4G). These results indicated that atRA could impair the stemness of MEPM cells and inhibit osteogenic differentiation.

Interestingly, we also found that when MEPM cells were separated from atRA-induced cleft palates, their LC3-II levels decreased, and their osteogenic-differentiation potency was weakened (Fig. 4A). Alizarin Red staining and RT-qPCR for osteogenic-related genes, ALP and Cbfα-1, confirmed these findings (Fig. 4G).

#### atRA inhibits the PTEN/Akt/mTOR autophagy signaling pathway in MEPM cells

We demonstrated that atRA regulated the stemness of MEPM cells and osteogenic differentiation through perturbation of autophagy in undifferentiated MEPM cells. We also verified that MEPM cells maintained pluripotency through the PTEN/Akt/mTOR autophagy signaling pathway. This raised the possibility for atRA regulation of autophagy in undifferentiated MEPM cells through inhibition of the PTEN/Akt/mTOR autophagy signaling pathway.Thus, we examined the signaling proteins, including pan- and phosphorylated-PTEN, Akt, and mTOR, that were treated with atRA doses of 5, 7, 10, and 15 μmol/mL for 24 h. Western blot showed that phospho-PTEN expression levels were upregulated upon atRA treatment at1, 3, 5, 7, 10, 15, and 20μmol/mL, while total PTEN protein expression was sharply downregulated. Consistent with altered PTEN levels, the downstream molecules of Akt and mTOR phosphorylation-two pivotal inhibitors of autophagy were upregulated without a concomitant effect on Akt and mTOR total protein expression (Fig. 4H). Collectively, these results demonstrated that atRA inhibited the PTEN/Akt/mTOR autophagy signaling pathway in MEPM cells.

#### Inhibition of PTEN/Akt/mTOR-mediated autophagy in atRA-induced cleft palate

At E10.5, pregnant mice were treated by oral gavage with atRA at 70 mg/kg according to our previous work^42^. All embryos showed cleft palate, and we observed litter death and other malformations; there was no cleft palate in normal groups (Table 3). atRA-induced E18.5 embryo groups showed a wider, complete secondary cleft palate compared to normal groups for failed elevated palates (Fig. 5A). Immunohistochemistry stainingof LC3-II showed that from E12.5 to E15.5, LC3-II levels decreased in atRA-induced groups compared with normal groups (Fig. 5B). These findings were confirmed by Western blot, in which the protein ratio of LC3-II to LC3-I was greatly decreased. Moreover, the levels of the autophagy adaptor protein p62 increased in the atRA-treated groups (Fig. 5C). These results were further verified by TEM: from E13.5 to E17.5, palatal shelves in the normal groups showed large numbers of autophagosomes, while in atRA-induced groups, autophagosome numbers were reduced (Fig. 5D). These results showed that atRA induced cleft palate by impairing autophagy, confirming that high basal levels of autophagy are characteristic of undifferentiated MEPM cells.

**Table 3.** Comparison of the incidence of cleft palate between atRA-induced and normal fetus.

|  | Dam | Total Fetus | Cleft Palate | Death | Ratio(%) |
| --- | --- | --- | --- | --- | --- |
|  | n | n | n | n |  |
| atRA-induced | 15 | 198 | 188 | 10 | 100% |
| Normal | 15 | 182 | 0 | 0 | 0 |

**Fig. 5.**
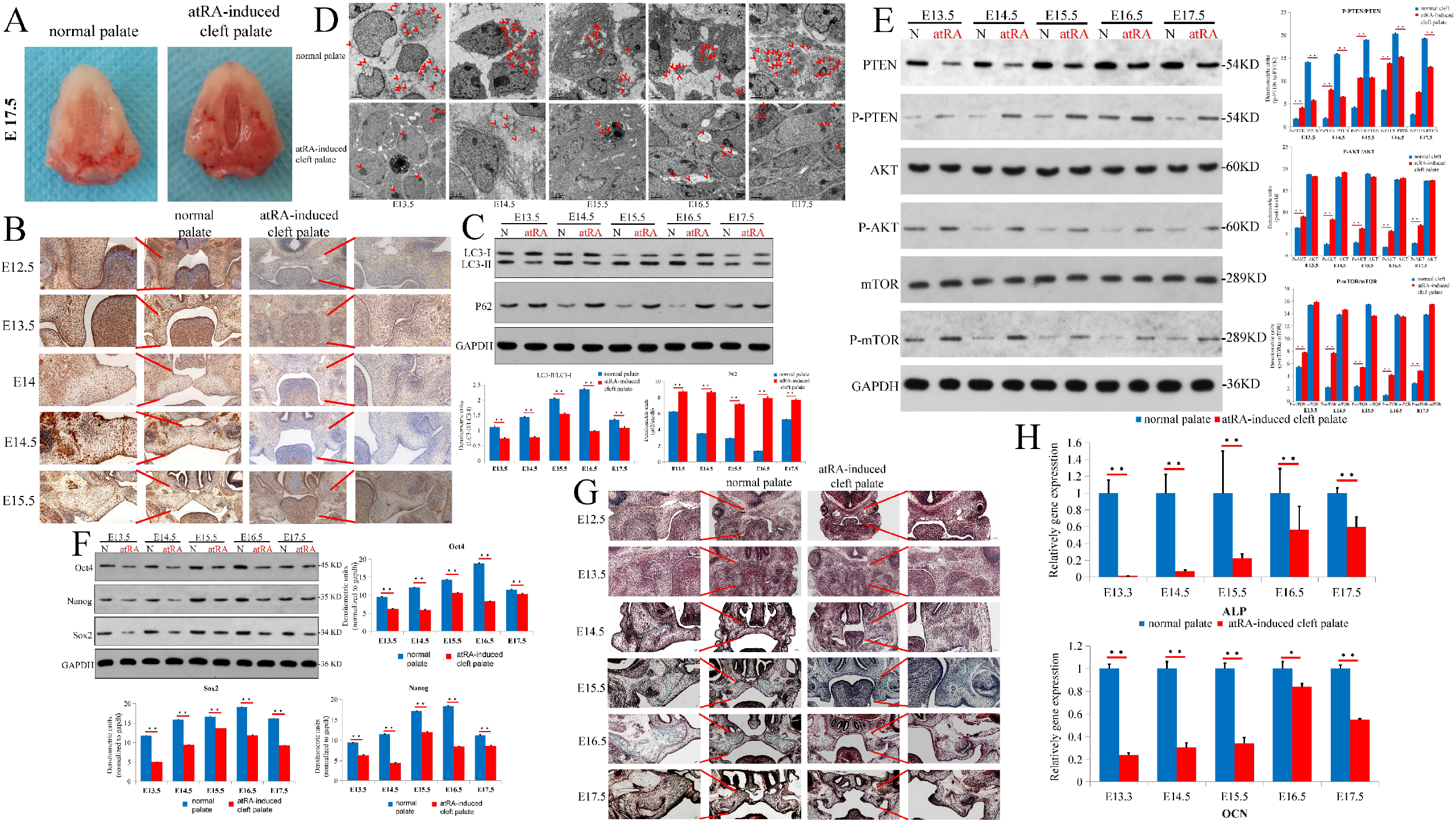
Characteristics of autophagy and ossification of palatal shelves in the atRA-induced cleft palate embryos. (A) Comparison of palatal general morphology between the normal palate and RA-induced cleft palate. (B) Immunohistochemical staining of LC3-II of palatal shelves from E12.5 to E17.5. Scale bar: 50 μm. (C) Results of Western blot analysis showing expressions of LC3-II/I and p62 proteins between normal and atRA-induced cleft palate from E12.5 to E17.5. (D) Transmission electron microscopy (TEM) image showing a few autophagosomes in the atRA-induced groups compared with normal groups from E12.5 to E17.5. (E) atRA blocked PTEN/Akt/mTOR autophagy signaling in atRA-induced cleft palate, as shown in Western blot analysis of protein levels of phosphorylated and non-phosphorylated PTEN, Akt, and mTOR from E12.5 to E17.5. (F) Western blot analysis of Oct4, Nanog, and Sox2 protein levels between atRA-induced and normal groups. (G) Goldner’s trichrome staining for palatal regions in the normal palate and atRA-induced cleft palate from E12.5 to E17.5. Mineralized bone: green; osteoid: orange/red; chondrocyte: purple; nuclei: blue/gray; cytoplasm: red/pink. Scale bar: 100 and 200μm. (H) Results of RT-qPCR showing upregulation of ALP and OCN in normal palate compared to the atRA-induced cleft palate. N = 3; *P < 0.05; **P < 0.01.

Having shown that atRA could block the PTEN/Akt/mTOR autophagy signaling pathway of MEPM cells *in vitro*, we also wondered whether this pathway was blocked in atRA-induced cleft palate. Western blot showed that expression levels of total PTEN were downregulated, while phosphorylation of PTEN, Akt, and mTOR was upregulated in palatal shelves from E13.5 to E17.5 in atRA-induced cleft palate relative to normal palate (Fig. 5E). These results were in accordance with the findings for MEPM cells cultured *in vitro*.

Having shown that atRA impairment of MEPM cells autophagy *in vitro* could affect cell stemness, we next analyzed the protein expression of stemness markers, including Oct4, Nanog, and Sox2, in atRA-induced cleft palate. Western blot results showed that from E13.5 to E17.5, expression of Oct4, Nanog, and Sox2 was downregulated in atRA-induced groups compared with normal groups (Fig. 5F). This result verified that stemness was impaired in atRA-induced cleft palate.

### Palatal-shelf ossification is weakened in atRA-induced cleft palate

Having shown that stemness was impaired in atRA-induced cleft palate, we next examined whether atRA also reduced the ossification of palatal shelves in this abnormality. Using Goldner’s trichrome staining to identify osteoidmineralized matrix, we found that ossification of palatal shelves from E12.5 to E17.5 was lower in the anterior regions of developing palates in atRA-induced groups than in normal groups (Fig. 5G). These findings were consistent with the RT-qPCR results, such that the osteogenic differentiation-related genes, ALP and OCN, were expressed at much lower levels in atRA-induced groups compared with normal groups (Fig. 5H). These results indicated that atRA indeed disturbed the ossification of palatal shelves to cause cleft palate.

### Differentially expressed microRNAs in atRA-induced mouse cleft palate orofacial tissues

The pleiotropic effects of atRA depend on two types of receptors—retinoic acid receptors (RARs) and retinoic X receptors (RXRs)—belonging to the nuclear hormone receptor superfamily^43,44^. We also found via IF that when MEPM cells were treated with atRA (10μmol/mL), their RARs were mainly located in the nucleus (Fig. 1D).

The PTEN/Akt/mTOR autophagy signaling pathway was located in the cytoplasmic membrane related to intracellular signaling pathways, but how atRA targets this pathway is unknown. MicroRNAs (miRNAs) reportedly act as regulatory signals for maintaining pluripotency, self-renewal, and differentiation of mesenchymal stem cells (MSCs)^45^. Additionally, microRNAs have emerged as important regulators of diverse physiological and pathological processes, with recent studies revealing several microRNAs that regulate MSC osteogenesis^46–49^. Therefore, we tested whether atRA targets the PTEN/Akt/mTOR autophagy signaling pathway via miRNAs.

To explore the participating miRNAs, we applied gene chip technology to the control and atRA treated groups on E14.5 to search for differentially expressed miRNAs. We detected 1126 microRNAs in all six fetal orofacial tissue samples by gene chip. Among them, 39 microRNAs were significantly upregulated, and 19 microRNAs were significantly downregulated in the atRA treated group. Heat map–based unsupervised hierarchical clustering analysis was performed with differentially expressed miRNAs (Fig. 6A). Among the differentially expressed microRNAs, the miR-106a-5p was the most significantly upregulated between the atRA-induced and normal groups (P = 0.0031, Fold change = 8.925) (Fig. 6B). Furthermore, we quantified miR-106a-5p levels in the palatal shelf between atRA-induced and normal groups from E13.5 to E17.5d using RT-qPCR. We found that miR-106a-5p levels were significantly increased between the atRA-induced and normal groups (Fig. 6C). At the same time, miR-106a-5p expression was detected in cultured MEPM cells that were treated with atRA; when exposed to atRA at different doses, miR-106a-5p expression increased with the increasing dose (Fig. 6D), thus verifying the results in the palatal shelf between the atRA-induced and normal groups. These results showed that atRA could induce miR-106a-5p expression in MEPM cells.

**Fig. 6.**
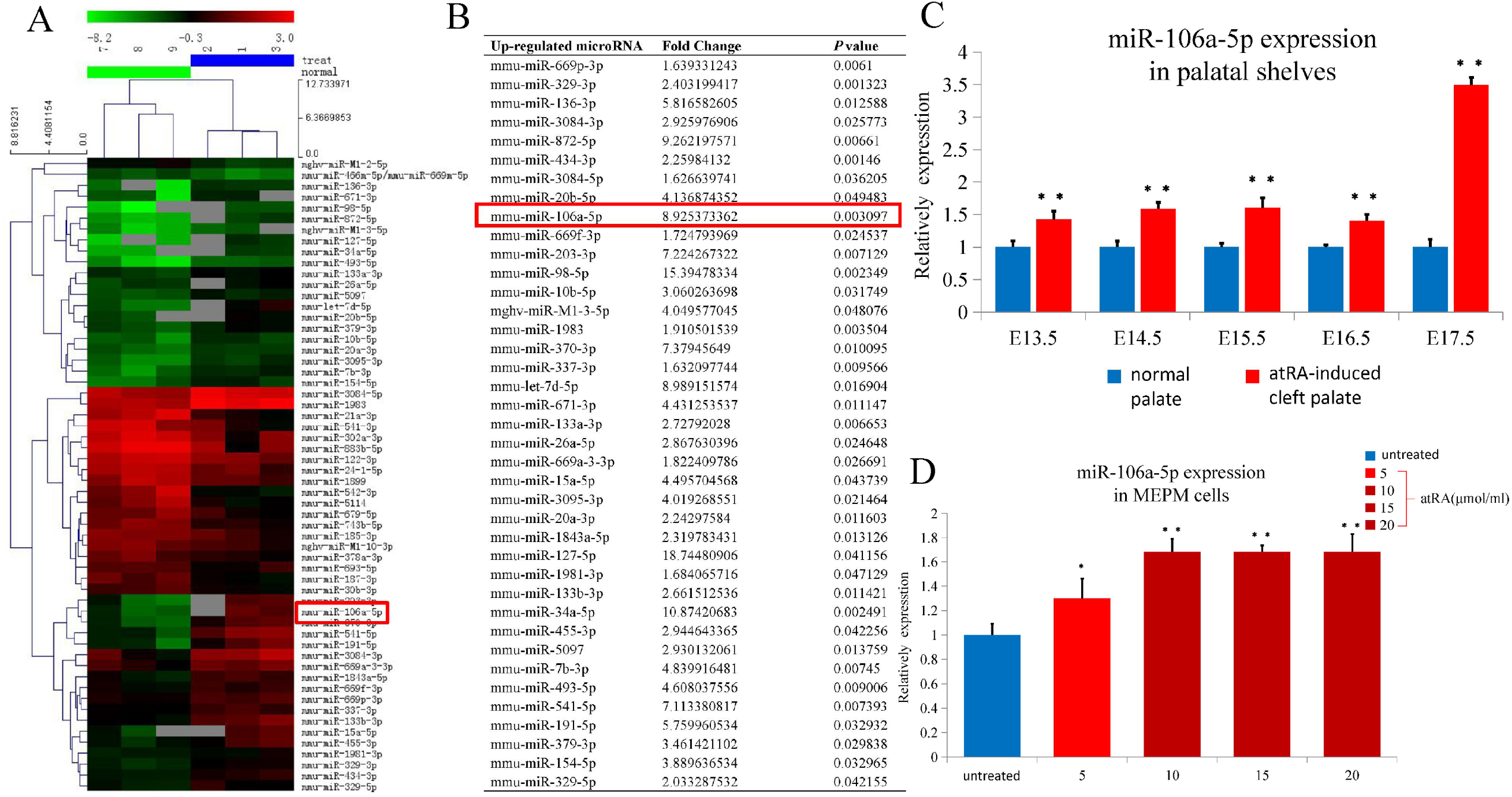
miRNA-106a-5p was significantly upregulated in atRA-induced cleft palate and atRA-treated MEPM cells. (A) Hierarchical clustering for differentially expressed miRNAs in atRA-induced cleft palate compared to the normal palate at E14.5. (B) Upregulated miRNAs in atRA-induced cleft palate compared to the normal palate at E14.5. (C) Results of RT-qPCR showing upregulation of miRNA-106a-5p in normal palate compared to atRA-induced cleft palate from E13.5-E17.5. (D) Results of RT-qPCR showing upregulation of miRNA-106a-5p in atRA-treated MEPM cells compared to untreated. N = 3; *P < 0.05; **P < 0.01.

### miR-106a-5p directly targets the 3’-UTR of PTEN mRNA to regulate the PTEN/Akt/mTOR autophagy signaling pathway

We wondered how miR-106a-5p regulates the PTEN/Akt/mTOR pathway. As such, we searched TargetScan (http://www.targetscan.org/vert_71/) to identify miR-106a-5p potential binding sites. We identified that the 3’UTR of PTEN matches the sequence of miR-106a-5p (Fig. 7A). Furthermore, to verify these predicted binding sites, we developed a dual-luciferase reporter system containing the wild type miR-106a-5p binding site and its mutant sequence that was co-transfected with either the miR-106a-5p over expression plasmid or the negative control along with a Renilla luciferase reporter system as a normalizer. The sequences of Pten-3’UTR-MT and Pten-3’UTR-WT were listed in Table4. Luciferase activity of the Pten-MT and miR-106a-5p mimic co-transfection group had no change compared to the negative control group (P <0.05); however, the luciferase activity of the Pten-WT and miRNA-106a-5p co-transfection group was significantly lower than the negative control group (P <0.05; Fig. 7B). These results confirmed that PTEN mRNA was a direct target of miR-106a-5p.

**Table 4.**
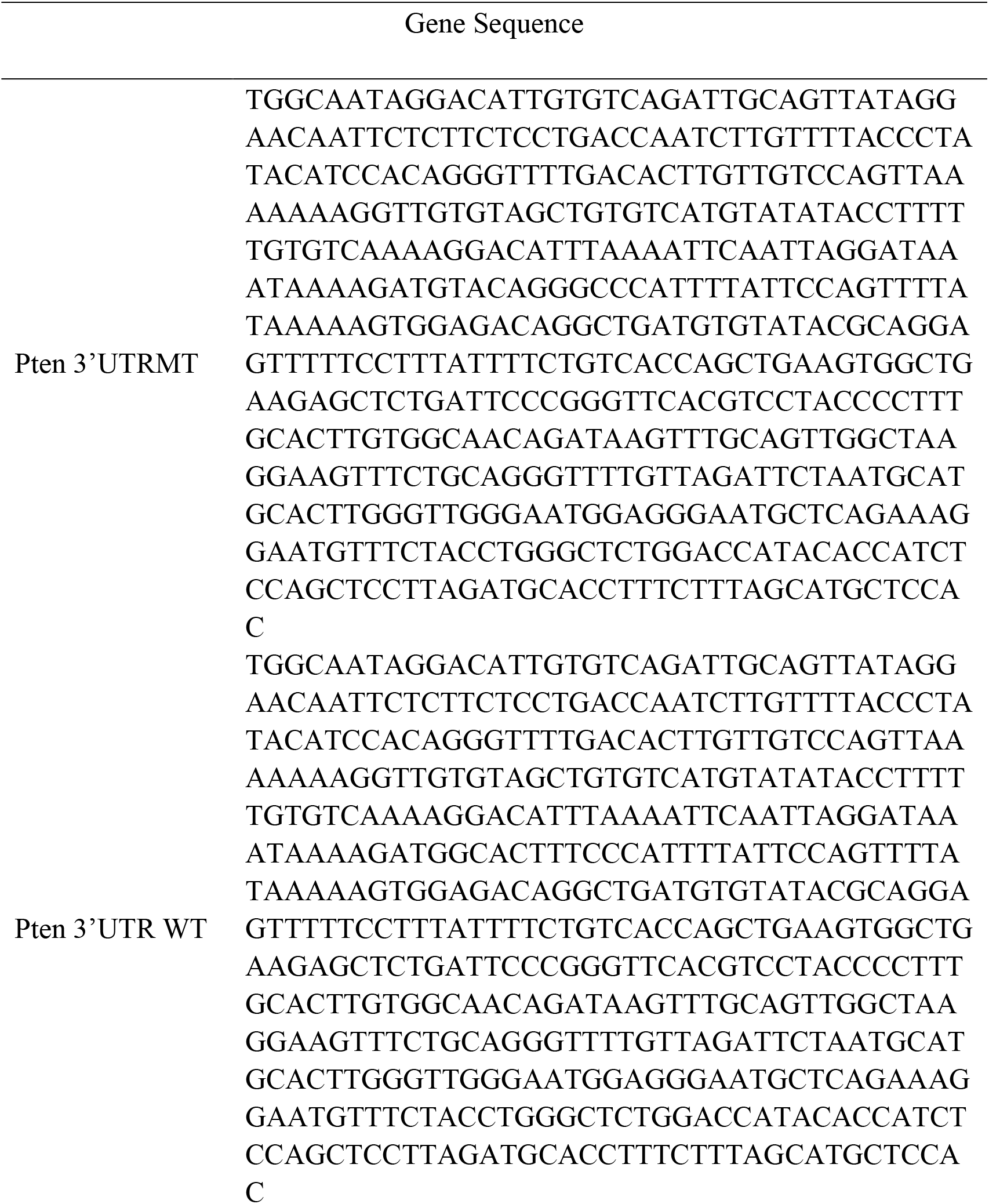
The sequence of Pten-3’UTR-MT and Pten-3-3’UTR-WT.

**Fig. 7.**
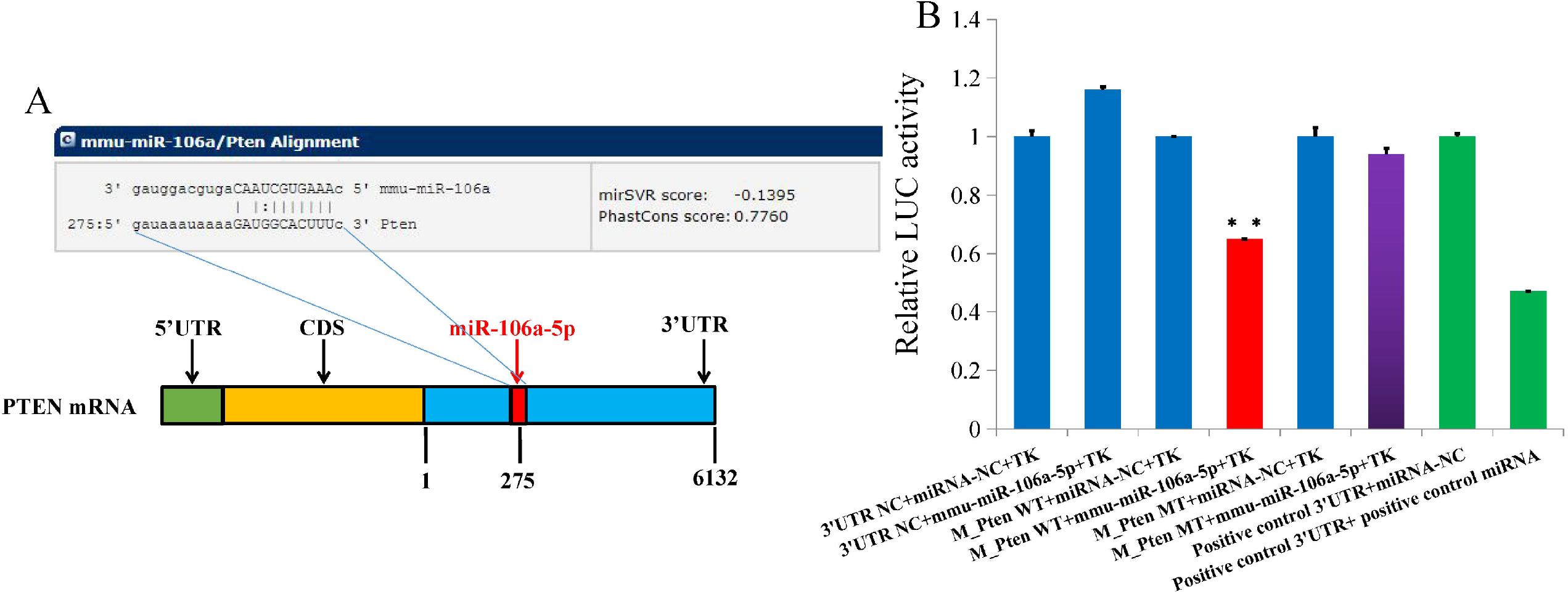
PTEN was a direct target of miRNA-106a-5p. (A) Potential miRNA-106a-5p binding site at the 3’UTR of PTEN mRNA was computationally predicted by TargetScan. (B) Luciferase activity was analyzed in cells co-transfected with miRNA-106a-5p mimics or negative control cells with pGL3-Pten 3’UTR-MT or pGL3-Pten 3’UTR-WT. The luciferases activity of group Pten-3’UTR-WT and miR-106a-5p-mimics co-transfection was significantly lower than in the Pten-NC and miR-106a-5p-NC group (**P<0.01). Group Pten-3’UTR-MT was used as control.There was no statistically significant difference from the Pten-MT and miR-106a-5p-mimics co-transfection group (P>0.05). N = 3.

### miR-106a-5p is upregulated in exosomes secreted by atRA-treated MEPM cells

TEM showed that MEPM cells secreted exosomes that were either cup-shaped or round morphologically (Fig. 8A). Nanosight analysis showed that the size of most exosomes secreted by MEPM cells was 82.13 nm (Fig. 8B). Furthermore, Western blot analyses showed that MEPM cells secreted exosomes that were positive for exosomal markers, including CD9, CD63, HSP70, and TSG101 (Fig. 8C). Furthermore, the expression of miR-106a-5p in exosomes secreted from atRA (10 μmol/mL)–treated and –untreated MEPM cells was examined with RT-qPCR. The results showed that miR-106a-5p was greatly upregulated in exosomes secreted by atRA-treated (10 μmol/mL) versus untreated MEPM cells (Fig. 8D).

**Fig. 8.**
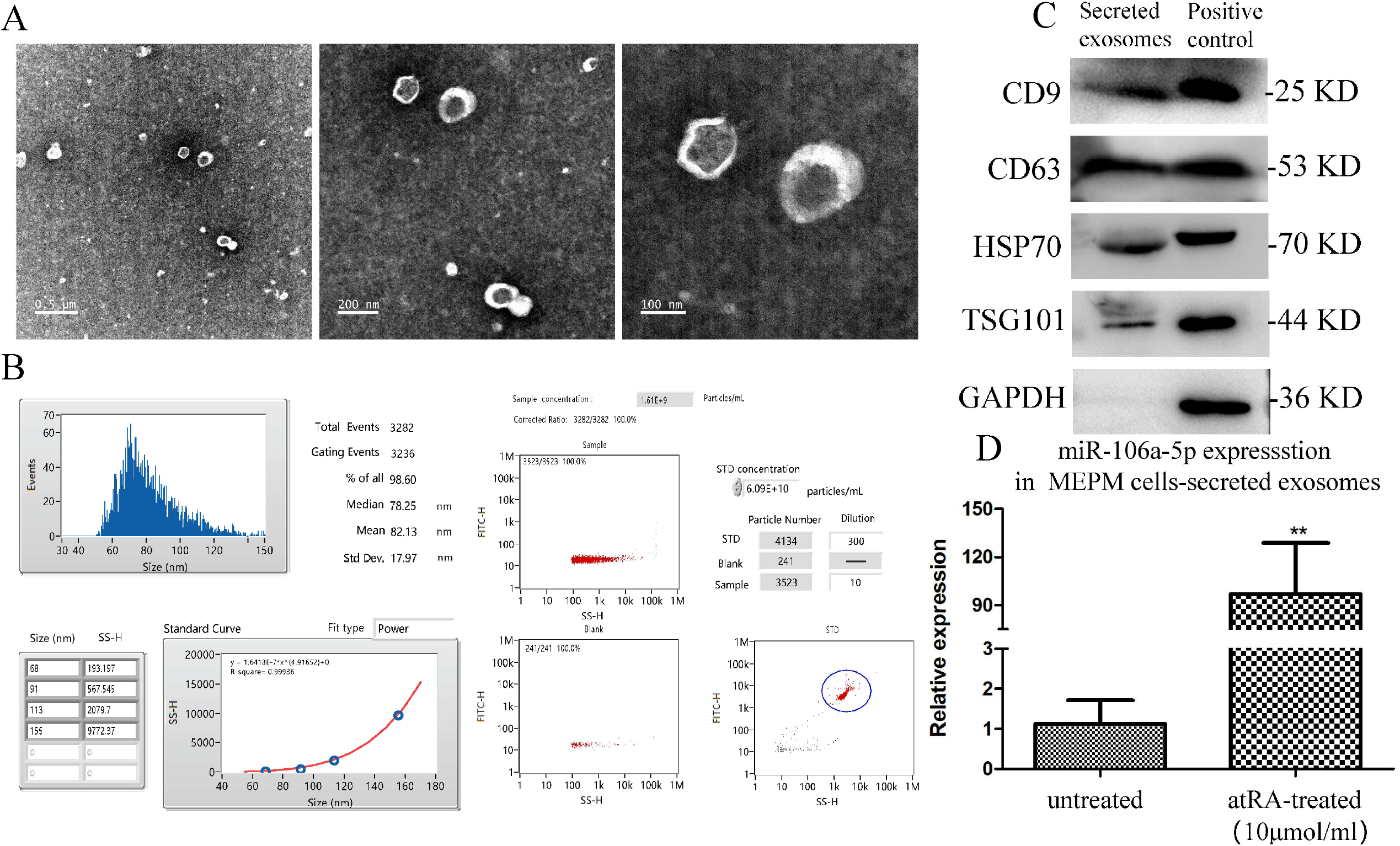
miR-106a-5p was upregulated in the atRA-treated MEPM cells secreted exosomes. (A) Morphology of exosomes identified by transmission electron microscopy (TEM). Scale bar, 200 nm. (B) Characteristics of atRA treated-MEPM cells secreted exosomes, NanoSight particle-size analysis of exosomes. (C) Expression of exosomal markers including CD63, CD9, CD81, and HSP70 measured using Western blot. (D) Results of RT-qPCR showed miR-106a-5p was upregulated in atRA-treated MEPM cells-secreted exosomes compared to untreated. N = 3; *P < 0.05; **P < 0.01.

These data indicated that exosomal miR-106a-5pwas a potential marker for cleft palate. Collectively, we concluded that atRA inhibited MEPM cell autophagy and osteogenic differentiation through miR-106a-5p targeting of PTEN and suppression of the PTEN/Akt/mTOR autophagy signaling pathway to ultimately cause cleft palate.

### miR-106a-5p inhibits the autophagy, stemness and osteogenic differentiation of MEPM cells through the PTEN/Akt/mTOR autophagy signaling pathway

As we had confirmed that atRA could upregulate the expression of miR-106a-5p, we also questioned whether miR-106a-5p could impair autophagy, stemness and osteogenic differentiation through the PTEN/Akt/mTOR autophagy signaling pathway. Therefore, we observed the expression of autophagy, stemness markers, including Oct4, Nanog, and Sox2; osteogenic differentiation; and the PTEN/Akt/mTOR autophagy signaling pathway of MEPM cells after transfection with miR-106a-5p. Western blot results showed that protein ratio of LC3-II to LC3-I was greatly decreased, the levels of the autophagy adaptor protein p62 increased after transfection with miR-106a-5p (Fig. 9A). Western blot results also showed the protein expression of Oct4, Nanog, and Sox2 was downregulated after transfection with miR-106a-5p (Fig. 9B), while after transfection with miR-106a-5p inhibitor, protein expression of these three markers was upregulated. Next, we explored the osteogenic-differentiation potency of MEPM cells transfected with miR-106a-5p. After 4 weeks of osteogenic induction, Alizarin Red staining showed a reduction in mineralized nodules after transfection with miR-106a-5p, while after transfectionwith miR-106a-5p inhibitor, mineralized nodules were increased (Fig. 9C). This was further verified by RT-qPCR analysis; osteogenic-related genes ALP and OCN showed a significant decrease in mRNA levels when cells were transfected with miR-106a-5p. Conversely, when transfected with miR-106a-5p inhibitor, expression of ALP and OCN was upregulated (Fig. 9D). These results indicated that miR-106a-5p could impair the autophagy, stemness of MEPM cells and inhibit osteogenic differentiation.

**Fig.9.**
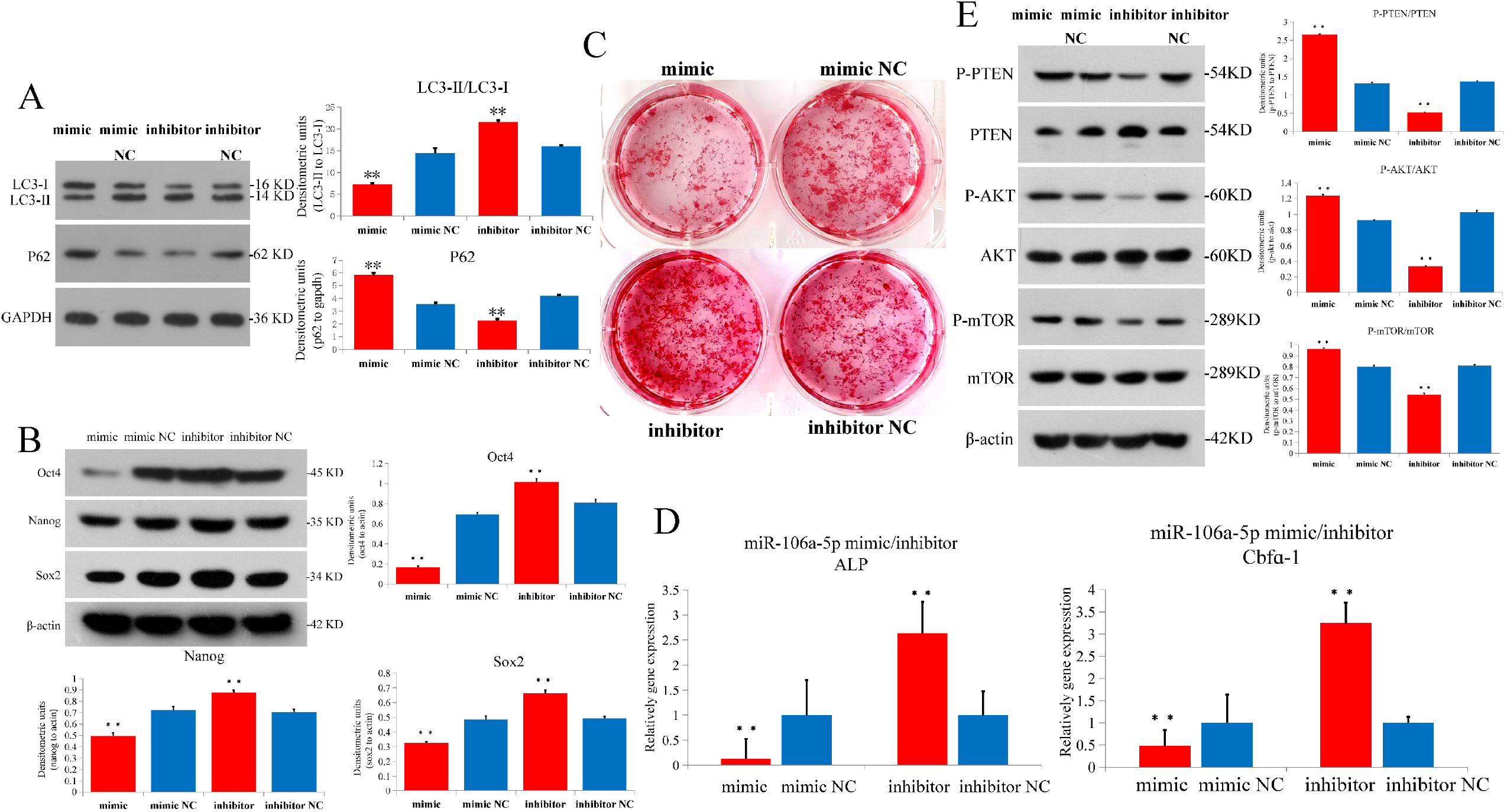
Characteristics of MEPM cells treated with miR-106a-5p. (A) Results of Western blot analysis showing expressions of LC3-II/I and p62 proteins after transfected with miR-106a-5p and inhibitor of miR-106a-5p. (B) Western blot analysis of Oct4, Nanog, and Sox2 protein levels in MEPM cells transfected with miR-106a-5p and inhibitor of miR-106a-5p. (C) After 4 weeks of osteogenic induction, Alizarin Red staining (ARS) showed weak calcium deposition when cells were transfected with miR-106a-5p (upper left panel), but increased calcium deposition when cells were transfected with miR-106a-5p inhibitor (bottom left panel). Scale bar: 50μm. (D) Results of RT-qPCR showing downregulation of ALP and OCN in MEPM cells transfected with miR-106a-5p and upregulation there of in MEPM cells transfected with miR-106a-5p inhibitor. (E) miR-106a-5p blocked the PTEN/Akt/mTOR autophagy signaling pathway, as shown by Western blot analysis of phosphorylated and nonphosphorylated PTEN, Akt, and mTOR protein levels. N = 3; *P < 0.05; **P < 0.01.

**Fig.10.**
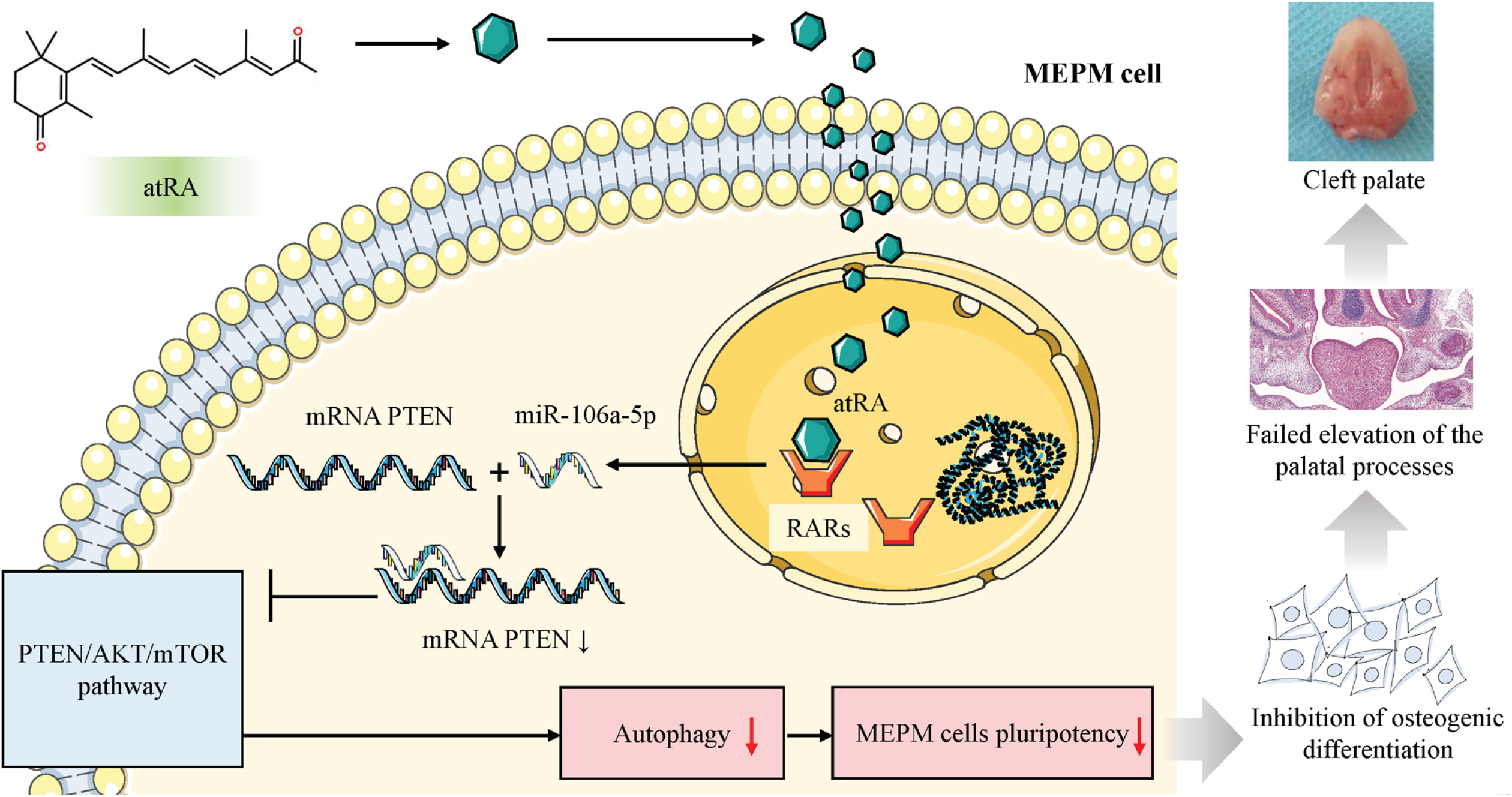
Schematic illustration of the PTEN/AKT/mTOR autophagy signaling pathway after atRA exposure in developing cleft palates.

Furthermore, we analyzed the PTEN/Akt/mTOR autophagy signaling pathway of MEPM cells transfected with miR-106a-5p. Western blot showed that after transfection with miR-106a-5p, the expression level of total PTEN was downregulated, while phosphorylation of PTEN, Akt, and mTOR was upregulated (Fig. 9E). After transfection with miR-106a-5p inhibitor, the expression level of total PTEN was upregulated, while phosphorylation of PTEN, Akt, and mTOR was downregulated (Fig. 9E). These results confirmed that miR-106a-5p could inhibit the PTEN/Akt/mTOR autophagy signaling pathway, which was consistent with the effects of atRA in the PTEN/Akt/mTOR autophagy signaling pathway.

## Discussion

Although atRA-induced cleft palate has been previously demonstrated^21^, the pathogenic mechanism for cleft palate caused by excess atRA exposure remains unclear. The cellular and molecular mechanisms for MEPM cell proliferation, and apoptosis and extracellular matrix production have been widely studied^21,50–54^; however, these studies do not fully explain the mechanism of atRA-induced cleft palate. It has been demonstrated that atRA exposure can permanently impede palatal shelf elevation, which is considered the main reason for the 100% incidence of cleft palates when embryonic mice are exposed to atRA on E10.5d^55,56^, and 90% of human cleft palate cases might attribute to the failure of the elevation process of palatal shelves^3,57^. It is thus necessary to fully elucidate the mechanism for vertical movement of the palatal shelves of atRA-induced cleft palate. We believe that atRA exposure can change the vertical movement of the palatal shelves at least two ways: (1) the decrease in proliferation of MEPM cells and increase in apoptosis that results in palatal shelves losing the capability of vertical growth; and (2) the osteogenic differentiation of MEPM cells is reduced, possibly obstructing the elevation of palatal shelves. Accordingly, osteogenic differentiation of MEPM cells may play a more important role in palatal-shelf elevation. Since the proliferation of MEPM cells must be based on osteogenic differentiation, the elevation of palatal shelves can be completed, similar to the lever principle. Once the osteogenic differentiation of MEPM cells is inhibited, even if the proliferation of MEPM cells is normal even increases, the palatal shelves could collapse and cause cleft palate.

Most MEPM cells (90%) derive from cranial neural crest cells (CNCCs)^58,59^; however, whether MEPM cells are still stem cells and possess multi-lineage differentiation potential is unclear. In this study, we verified that MEPM cells were ectomesenchymal stem cells and showed strong osteogenic differentiation potential. The perturbation of stemness can affect proliferation capacities and differentiation potential. atRA is often considered a differentiation agent in leukemia and neural differentiation^60–63^. Therefore, atRA can affect stemness and pluripotency and impede the differentiation potential of stem cells.Accordingly, atRA could affect the stemness and pluripotency of MEPM cells, resulting in abnormal MEPM proliferation and differentiation, which can cause cleft palate.

The hard palate is a common bony partition between the oral and nasal cavity and comprises two-thirds of the palate^64^. Therefore, the ossification of palate shelves plays a critical role during the development of palatal shelves^65–67^. We found in atRA-induced cleft palates that the ossification of palatal shelves was much weaker *in vivo* from E12.5 to E17.5 than in samples associated with normal palate. We also found the same downregulated of the stemness markers, such as Oct4, Nanog, and Sox2, in atRA-induced cleft palates. Palatal-shelf mineralization is achieved by in tramembranous ossification where MEPM cells could directly differentiate into osteoblasts. To verify whether atRA could affect MEPM cells stemness and osteogenic differentiation, we found that stemness markers, such as Oct4, Nanog, and Sox2, also showed less expression when exposed to atRA. We further induced MEPM cells that were exposed to different doses of atRA for 24h to osteogenic differentiation for 4 weeks, and found that when exposed to a dose of atRA up to 10 μmol/mL, the osteogenic differentiation sharply decreased. This verified that atRA could disturb MEPM cells stemness and osteogenic differentiation. These results are consistent with several studies (see Cohen-Tanugi and Forest, 1998; Skillington *et al*., 2002; Wang *et al*., 2008; Chen *et al*., 2010; Wang *et al*., 2016)^68–72^. We also induced MEPM cells isolated from palatal shelves from atRA-induced cleft palate to osteogenic differentiation for 4 weeks. Interestingly, the osteogenic differentiation of MEPM cells from atRA-induced cleft palates were also weaker than MEPM cells from normal palates. We did not determine whether atRA could affect adipogenic and chondrogenic differentiation, but there are many other studies that show that atRA could block adipogenic differentiation by MEPM cells^73–75^ and chondrogenic differentiation^76–80^. Our results showed that atRA regulated osteogenic differentiation through the disruption of MEPM cell stemness and pluripotency.

Several studies report that high levels of autophagy are critical for stem cell survival when subjected to harsh conditions^81^. In this study, we observed a high density of autophagosomes in undifferentiated MEPM cells; however, upon differentiation into osteogenic cells, the autophagosomes disappeared. We also found when the MEPM cells differentiated into adipogenic and chondrogenic cells, the autophagosomes also disappeared (data not shown). This suggests that in an undifferentiated state, MEPM cells have a high autophagosome concentration. Therefore, the level of autophagy may be considered as a hallmark of undifferentiated MEPM cells. By contrast, we found that when MEPM cells were treated with atRA, the levels of autophagy decreased, and when the dose of atRA reached 10μmol/mL, the autophagosomes disappeared. In this way, our study verified that atRA can inhibit MEPM cells stemness and pluripotency through disturbing the autophagy.

The PI3K/Akt/mTOR signaling pathway plays a central role in autophagy by regulating growth, motility, protein synthesis, metabolism, survival, and death from various stimuli^82,83^. While PTEN acts as a suppressor of PI3K and Akt, the activity of PTEN also can be negatively regulated by phosphorylation, resulting in the inactivation of PTEN^84^. In this study, PTEN phosphorylation was activated by osteogenic differentiation, leading to PI3K/Akt/mTOR signaling pathway activation. Furthermore, we found that when undifferentiated MEPM cells were treated with the PTEN inhibitor, VO-OHpic trihydrate, which was capable of blocking the PTEN/AKT/mTOR signaling pathway (Fig. 3C), the autophagosome numbers and osteogenic differentiation potential decreased. Taken together, we concluded that undifferentiated MEPM cells hold a high basal level of autophagy to maintain pluripotency through PTEN/AKT/mTOR signaling activation. Conversely, we found that when undifferentiated MEPM cells were treated with atRA, the PTEN was inactive, while the phosphorylated PTEN was active. Additionally, in atRA-induced cleft palate, we found that PTEN/AKT/mTOR signaling was blocked. Our results are consistent with previous work (Susana Masiá *et al*., 2007; Busada *et al*., 2015)^85,86^, as their studies show that atRA could utilize non-genomic pathways via PI3K/AKT/mTOR signaling to stimulate translation of mRNA. Thus, our results showed that atRA could block PTEN/AKT/mTOR autophagy signaling, which could result in cleft palate formation.

Retinoic acid receptors (RARs) belong to the nuclear hormone receptor superfamily^43,44^. Many studies show that RARs are not in the plasma membrane of SH-SY5Y, NIH3T3, and MEFs^85,87,88^. We confirmed that RARs were located in the nucleus, which were not observed in MEPM cell plasma membranes (Fig. 1D). However, PTEN/AKT/mTOR signaling occurs in the plasma membrane. Therefore, there is a missing link in how atRA affects PTEN/AKT/mTOR signaling. We speculate that miR-106a-5p may play an important role since miRNAs are key regulators in palatogenesis and cleft palate^89–91^, and because atRA exposure can induce the expression of different miRNAs^92–96^. In this study, we demonstrated that miR-106a-5p was significantly upregulated in atRA-exposed MEPM cells through the identification of differentially expressed microRNAs in orofacial tissues obtained at E14.5 following atRA exposure. Additionally, we found that miR-106a-5p was upregulated when MEPM cells were treated with atRA for 24h, verifying that atRA induced miR-106a-5p expression. Further, the dual-luciferase reporter system confirmed that PTEN mRNA was a direct target of miR-106a-5p, which inhibited PTEN expression to block PTEN/AKT/mTOR signaling and, ultimately, reduced the osteogenic differentiation of MEPM cells. Our results are consistent with previous studies demonstrating that miR-106a-5p upregulation results in the reduction of the osteogenic differentiation of mesenchymal stem cells^97–100^. Manochantr *et al*.^101^ found that during the osteogenic differentiation of mesenchymal stem cells derived from amnion, miR-106a is downregulated; however, anti-miR-106acouldpromote the osteogenic process.

In conclusion, our findings indicate that *in vitro* purified MEPM cells are ectomesenchymal stem cells that have the differential potential for osteogenic lineages; this pluripotency and differentiation potential is regulated by autophagy through PTEN/AKT/mTOR signaling. atRA disturbs the MEPM cell pluripotency through the inactivation of the PTEN/AKT/mTOR pathway via miR-106a-5p targeting PTEN mRNA and reduces MEPM cell osteogenic differentiation that results in the development of cleft palate. Our findings provide new insight for understanding the mechanism for cleft palate development. We identify miR-106a-5p as a biomarker for prenatal screening for cleft palate and as a new diagnostic and therapeutic target for cleft palate. Future studies are needed to replicate these findings in humans, including verifying whether miR-106a-5p is expressed in exosomes of cleft palate patient blood.

## Declarations

## Acknowledgments

We would like to thank LetPub (www.letpub.com) for providing linguistic assistance during the preparation of this manuscript.

## Availability of data and material

The authors declare that all the data supporting the findings of this study are available within the article and that no data sharing is applicable to this article.

## Authors’ contributions

LS conceived the study and experimental design and executed all the molecular and cellular experiments. YL, BZ and LY analyzed the data. BL and CZ were responsible for the literature review.JF and ST contributed to the conception of this manuscript. LS and ST drafted and revised the manuscript. All authors read and approved the final manuscript.

## Ethics approval

All animal studies were approved by the Laboratory Animal Ethical Committee of the Medical College of the Shantou University.

## Consent for publication

Not applicable.

## References

1 Meng, L., Bian, Z., Torensma, R. & Von den Hoff, J. W. Biological mechanisms in palatogenesis and cleft palate. Journal of dental research 88, 22–33, doi:10.1177/0022034508327868 (2009).

2 Funato, N., Nakamura, M. & Yanagisawa, H. Molecular basis of cleft palates in mice. World journal of biological chemistry 6, 121–138, doi:10.4331/wjbc.v6.i3.121 (2015).

3 Ferguson, M. W. Palate development. Development 103 Suppl, 41–60 (1988).

4 Meng, T. et al. Overexpression of mouse TTF-2 gene causes cleft palate. Journal of cellular and molecular medicine 16, 2362–2368, doi:10.1111/j.1582-4934.2012.01546.x (2012).

5 Wu, C. et al. Intra-amniotic transient transduction of the periderm with a viral vector encoding TGFbeta3 prevents cleft palate in Tgfbeta3(-/-) mouse embryos. Mol Ther 21, 8–17, doi:10.1038/mt.2012.135 (2013).

6 Murray, J. C. & Schutte, B. C. Cleft palate: players, pathways, and pursuits. Journal of Clinical Investigation 113, 1676–1678, doi:10.1172/jci200422154 (2004).

7 Khrapunov, S. M., Zima, V. L., Tiuleniev, V. I. & Berdishev, H. D. [Change in conformation of histones F 2a and F 2b in solutions of different ionic strength]. Ukrains’kyi biokhimichnyi zhurnal 47, 284–289 (1975).

8 Shuler, C. F. Programmed cell death and cell transformation in craniofacial development. Critical reviews in oral biology and medicine : an official publication of the American Association of Oral Biologists 6, 202–217 (1995).

9 Wilkie, A. O. & Morriss-Kay, G. M. Genetics of craniofacial development and malformation. Nature reviews. Genetics 2, 458–468, doi:10.1038/35076601 (2001).

10 Li, L., Shi, J. Y., Zhu, G. Q. & Shi, B. MiR-17-92 cluster regulates cell proliferation and collagen synthesis by targeting TGFB pathway in mouse palatal mesenchymal cells. Journal of cellular biochemistry 113, 1235–1244, doi:10.1002/jcb.23457 (2012).

11 Clagett-Dame, M. & DeLuca, H. F. The role of vitamin A in mammalian reproduction and embryonic development. Annu Rev Nutr 22, 347–381, doi:10.1146/annurev.nutr.22.010402.102745E (2002).

12 Dmetrichuk, J. M., Spencer, G. E. & Carlone, R. L. Retinoic acid-dependent attraction of adult spinal cord axons towards regenerating newt limb blastemas in vitro. Dev Biol 281, 112–120, doi:10.1016/j.ydbio.2005.02.019 (2005).

13 Wang, M., Huang, H. & Chen, Y. Smad2/3 is involved in growth inhibition of mouse embryonic palate mesenchymal cells induced by all-trans retinoic acid. Birth Defects Res A Clin Mol Teratol 85, 780–790, doi:10.1002/bdra.20588 (2009).

14 <abbott1989.pdf>.

15 Mizushima, N. & Levine, B. Autophagy in mammalian development and differentiation. Nat Cell Biol 12, 823–830, doi:10.1038/ncb0910-823 (2010).

16 Wu, X., Won, H. & Rubinsztein, D. C. Autophagy and mammalian development. Biochem Soc Trans 41, 1489–1494, doi:10.1042/BST20130185 (2013).

17 Komatsu, M. et al. Loss of autophagy in the central nervous system causes neurodegeneration in mice. Nature 441, 880–884, doi:10.1038/nature04723 (2006).

18 Lee, H. K. & Iwasaki, A. Autophagy and antiviral immunity. Curr Opin Immunol 20, 23–29, doi:10.1016/j.coi.2008.01.001 (2008).

19 Levine, B. & Kroemer, G. Autophagy in the pathogenesis of disease. Cell 132, 27–42, doi:10.1016/j.cell.2007.12.018 (2008).

20 Hu, X., Chen, Z., Mao, X. & Tang, S. Effects of phenytoin and Echinacea purpurea extract on proliferation and apoptosis of mouse embryonic palatal mesenchymal cells. Journal of cellular biochemistry 112, 1311–1317, doi:10.1002/jcb.23044 (2011).

21 Hu, X., Gao, J., Liao, Y., Tang, S. & Lu, F. Retinoic acid alters the proliferation and survival of the epithelium and mesenchyme and suppresses Wnt/beta-catenin signaling in developing cleft palate. Cell death & disease 4, e898, doi:10.1038/cddis.2013.424 (2013).

22 Baddoo, M. et al. Characterization of mesenchymal stem cells isolated from murine bone marrow by negative selection. Journal of cellular biochemistry 89, 1235–1249, doi:10.1002/jcb.10594 (2003).

23 Wu, M. et al. Persistent expression of Pax3 in the neural crest causes cleft palate and defective osteogenesis in mice. The Journal of clinical investigation 118, 2076–2087, doi:10.1172/JCI33715 (2008).

24 Yang, B., Guo, H., Zhang, Y., Dong, S. & Ying, D. The microRNA expression profiles of mouse mesenchymal stem cell during chondrogenic differentiation. BMB reports 44, 28–33, doi:10.5483/BMBRep.2011.44.1.28 (2011).

25 Liu, A. R. et al. Activation of canonical wnt pathway promotes differentiation of mouse bone marrow-derived MSCs into type II alveolar epithelial cells, confers resistance to oxidative stress, and promotes their migration to injured lung tissue in vitro. Journal of cellular physiology 228, 1270–1283, doi:10.1002/jcp.24282 (2013).

26 Chen, Y. et al. Mesenchymal-like stem cells derived from human parthenogenetic embryonic stem cells. Stem cells and development 21, 143–151, doi:10.1089/scd.2010.0585 (2012).

27 Hwang, N. S. et al. In vivo commitment and functional tissue regeneration using human embryonic stem cell-derived mesenchymal cells. Proceedings of the National Academy of Sciences of the United States of America 105, 20641–20646, doi:10.1073/pnas.0809680106 (2008).

28 Liu, Q. et al. A comparative study of proliferation and osteogenic differentiation of adipose-derived stem cells on akermanite and beta-TCP ceramics. Blomazerlals 29, 4792–4799, doi:10.1016/j.biomaterials.2008.08.039 (2008).

29 Gregory, C. A., Gunn, W. G., Peister, A. & Prockop, D. J. An Alizarin red-based assay of mineralization by adherent cells in culture: comparison with cetylpyridinium chloride extraction. Analytical biochemistry 329, 77–84, doi:10.1016/j.ab.2004.02.002 (2004).

30 Mao, G. et al. Exosomal miR-95-5p regulates chondrogenesis and cartilage degradation via histone deacetylase 2/8. Journal of cellular and molecular medicine 22, 5354–5366, doi:10.1111/jcmm.13808 (2018).

31 Shi, L. et al. Mouse embryonic palatal mesenchymal cells maintain stemness through the PTEN-Akt-mTOR autophagic pathway. Stem Cell Res Ther 10, 217, doi:10.1186/s13287-019-1340-8 (2019).

32 Greening, D. W., Xu, R., Ji, H., Tauro, B. J. & Simpson, R. J. A protocol for exosome isolation and characterization: evaluation of ultracentrifugation, density-gradient separation, and immunoaffinity capture methods. Methods Mol Biol 1295, 179–209, doi:10.1007/978-1-4939-2550-6_15 (2015).

33 Chambers, I. et al. Functional expression cloning of Nanog, a pluripotency sustaining factor in embryonic stem cells. Cell 113, 643–655, doi:10.1016/s0092-8674(03)00392-1 (2003).

34 Boyer, L. A. et al. Core transcriptional regulatory circuitry in human embryonic stem cells. Cell 122, 947–956, doi:10.1016/j.cell.2005.08.020 (2005).

35 Mendez-Ferrer, S. et al. Mesenchymal and haematopoietic stem cells form a unique bone marrow niche. Nature 466, 829–834, doi:10.1038/nature09262 (2010).

36 Gazarian, K. G. & Ramirez-Garcia, L. R. Human Deciduous Teeth Stem Cells (SHED) Display Neural Crest Signature Characters. PloS one 12, e0170321, doi:10.1371/journal.pone.0170321 (2017).

37 Oliver, L., Hue, E., Priault, M. & Vallette, F. M. Basal autophagy decreased during the differentiation of human adult mesenchymal stem cells. Stem cells and development 21, 2779–2788, doi:10.1089/scd.2012.0124 (2012).

38 Vessoni, A. T., Muotri, A. R. & Okamoto, O. K. Autophagy in stem cell maintenance and differentiation. Stem cells and development 21, 513–520, doi:10.1089/scd.2011.0526 (2012).

39 Zhang, Q. et al. Autophagy Activation: A Novel Mechanism of Atorvastatin to Protect Mesenchymal Stem Cells from Hypoxia and Serum Deprivation via AMP-Activated Protein Kinase/Mammalian Target of Rapamycin Pathway. Stem cells and development 21, 1321–1332, doi:10.1089/scd.2011.0684 (2012).

40 Vucicevic, L. et al. Compound C induces protective autophagy in cancer cells through AMPK inhibition-independent blockade of Akt/mTOR pathway. Autophagy 7, 40–50, doi:10.4161/auto.7.1.13883 (2014).

41 Li, X. Y. et al. Triptolide Restores Autophagy to Alleviate Diabetic Renal Fibrosis through the miR-141-3p/PTEN/Akt/mTOR Pathway. Mol Ther Nucleic Acids 9, 48–56, doi:10.1016/j.omtn.2017.08.011 (2017).

42 Zhang, W. et al. MiR-106a-5p modulates apoptosis and metabonomics changes by TGF-beta/Smad signaling pathway in cleft palate. Exp Cell Res 386, 111734, doi:10.1016/j.yexcr.2019.111734 (2020).

43 Chambon, P. A decade of molecular biology of retinoic acid receptors. FASEB J 10, 940–954 (1996).

44 Bastien, J. & Rochette-Egly, C. Nuclear retinoid receptors and the transcription of retinoid-target genes. Gene 328, 1–16, doi:10.1016/j.gene.2003.12.005 (2004).

45 Wu, T. et al. miR-21 Modulates the Immunoregulatory Function of Bone Marrow Mesenchymal Stem Cells Through the PTEN/Akt/TGF-beta1 Pathway. Stem Cells 33, 3281–3290, doi:10.1002/stem.2081 (2015).

46 Kim, K. M. et al. miR-182 is a negative regulator of osteoblast proliferation, differentiation, and skeletogenesis through targeting FoxO1. J Bone Miner Res 27, 1669–1679, doi:10.1002/jbmr.1604 (2012).

47 Hassan, M. Q. et al. A network connecting Runx2, SATB2, and the miR-23a~27a~24-2 cluster regulates the osteoblast differentiation program. Proceedings of the National Academy of Sciences of the United States of America 107, 19879–19884, doi:10.1073/pnas.1007698107 (2010).

48 Eskildsen, T. et al. MicroRNA-138 regulates osteogenic differentiation of human stromal (mesenchymal) stem cells in vivo. Proceedings of the National Academy of Sciences of the United States of America 108, 6139–6144, doi:10.1073/pnas.1016758108 (2011).

49 Huang, J., Zhao, L., Xing, L. & Chen, D. MicroRNA-204 regulates Runx2 protein expression and mesenchymal progenitor cell differentiation. Stem Cells 28, 357–364, doi:10.1002/stem.288 (2010).

50 Cuervo, R., Valencia, C., Chandraratna, R. A. & Covarrubias, L. Programmed cell death is required for palate shelf fusion and is regulated by retinoic acid. Dev Biol 245, 145–156, doi:10.1006/dbio.2002.0620 (2002).

51 Kawaguchi, J., Mee, P. J. & Smith, A. G. Osteogenic and chondrogenic differentiation of embryonic stem cells in response to specific growth factors. Bone 36, 758–769, doi:10.1016/j.bone.2004.07.019 (2005).

52 Choi, J. W., Park, H. W., Kwon, Y. J. & Park, B. Y. Role of apoptosis in retinoic acid-induced cleft palate. J Craniofac Surg 22, 1567–1571, doi:10.1097/SCS.0b013e318208ba10 (2011).

53 Schroen, D. J. & Brinckerhoff, C. E. Nuclear hormone receptors inhibit matrix metalloproteinase (MMP) gene expression through diverse mechanisms. Gene Expr 6, 197–207 (1996).

54 Liu, S., Higashihori, N., Yahiro, K. & Moriyama, K. Retinoic acid inhibits histone methyltransferase Whsc1 during palatogenesis. Biochem Biophys Res Commun 458, 525–530, doi:10.1016/j.bbrc.2015.01.148 (2015).

55 Qin, F. et al. Metabolic characterization of all-trans-retinoic acid (ATRA)-induced craniofacial development of murine embryos using in vivo proton magnetic resonance spectroscopy. PloS one 9, e96010, doi:10.1371/journal.pone.0096010 (2014).

56 Shu, X., Shu, S., Zhai, Y., Zhu, L. & Ouyang, Z. Genome-Wide DNA Methylation Profile of Gene cis-Acting Element Methylations in All-trans Retinoic Acid-Induced Mouse Cleft Palate. DNA Cell Biol, doi:10.1089/dna.2018.4369 (2018).

57 Ferguson, M. W. The mechanism of palatal shelf elevation and the pathogenesis of cleft palate. Virchows Arch A Pathol Anat Histol 375, 97–113 (1977).

58 Iwata, J., Parada, C. & Chai, Y. The mechanism of TGF-beta signaling during palate development. Oral Dis 17, 733–744, doi:10.1111/j.1601-0825.2011.01806.x (2011).

59 Ito, Y. et al. Conditional inactivation of Tgfbr2 in cranial neural crest causes cleft palate and calvaria defects. Development 130, 5269–5280, doi:10.1242/dev.00708 (2003).

60 Mongan, N. P. & Gudas, L. J. Diverse actions of retinoid receptors in cancer prevention and treatment. Differentiation 75, 853–870, doi:10.1111/j.1432-0436.2007.00206.x (2007).

61 Angrisano, T. et al. Chromatin and DNA methylation dynamics during retinoic acid-induced RET gene transcriptional activation in neuroblastoma cells. Nucleic Acids Res 39, 1993–2006, doi:10.1093/nar/gkq864 (2011).

62 Janesick, A., Wu, S. C. & Blumberg, B. Retinoic acid signaling and neuronal differentiation. Cellular and molecular life sciences : CMLS 72, 1559–1576, doi:10.1007/s00018-014-1815-9 (2015).

63 Zhu, H. H. & Huang, X. J. Oral arsenic and retinoic acid for non-high-risk acute promyelocytic leukemia. N Engl J Med 371, 2239–2241, doi:10.1056/NEJMc1412035 (2014).

64 Okano, J., Udagawa, J. & Shiota, K. Roles of retinoic acid signaling in normal and abnormal development of the palate and tongue. Congenit Anom (Kyoto) 54, 69–76, doi:10.1111/cga.12049 (2014).

65 Wu, X., Liu, X. & Wang, S. Implantation of biomaterial as an efficient method to harvest mesenchymal stem cells. Experimental biology and medicine 236, 1477–1484, doi:10.1258/ebm.2011.011061 (2011).

66 Fu, X. et al. Identification of Osr2 Transcriptional Target Genes in Palate Development. Journal of dental research 96, 1451–1458, doi:10.1177/0022034517719749 (2017).

67 Jia, S. et al. Small-molecule Wnt agonists correct cleft palates in Pax9 mutant mice in utero. Development 144, 3819–3828, doi:10.1242/dev.157750 (2017).

68 Cohen-Tanugi, A. & Forest, N. Retinoic acid suppresses the osteogenic differentiation capacity of murine osteoblast-like 3/A/1D-1M cell cultures. Differentiation 63, 115–123, doi:10.1046/j.1432-0436.1998.6330115.x (1998).

69 Skillington, J., Choy, L. & Derynck, R. Bone morphogenetic protein and retinoic acid signaling cooperate to induce osteoblast differentiation of preadipocytes. J Cell Biol 159, 135–146, doi:10.1083/jcb.200204060 (2002).

70 Wang, A., Ding, X., Sheng, S. & Yao, Z. Retinoic acid inhibits osteogenic differentiation of rat bone marrow stromal cells. Biochem Biophys Res Commun 375, 435–439, doi:10.1016/j.bbrc.2008.08.036 (2008).

71 Chen, M., Huang, H. Z., Wang, M. & Wang, A. X. Retinoic acid inhibits osteogenic differentiation of mouse embryonic palate mesenchymal cells. Birth Defects Res A Clin Mol Teratol 88, 965–970, doi:10.1002/bdra.20723 (2010).

72 Wang, W. et al. All-Trans Retinoic Acid-Induced Craniofacial Malformation Model: A Prenatal and Postnatal Morphological Analysis. Cleft Palate Craniofac J 54, 391–399, doi:10.1597/15-271 (2017).

73 Wang, B., Fu, X., Zhu, M. J. & Du, M. Retinoic acid inhibits white adipogenesis by disrupting GADD45A-mediated Zfp423 DNA demethylation. J Mol Cell Biol 9, 338–349, doi:10.1093/jmcb/mjx026 (2017).

74 Berry, D. C. & Noy, N. All-trans-retinoic acid represses obesity and insulin resistance by activating both peroxisome proliferation-activated receptor beta/delta and retinoic acid receptor. Mol Cell Biol 29, 3286–3296, doi:10.1128/MCB.01742-08 (2009).

75 Berry, D. C., DeSantis, D., Soltanian, H., Croniger, C. M. & Noy, N. Retinoic acid upregulates preadipocyte genes to block adipogenesis and suppress diet-induced obesity. Diabetes 61, 1112–1121, doi:10.2337/db11-1620 (2012).

76 Yu, Z. & Xing, Y. All-trans retinoic acid inhibited chondrogenesis of mouse embryonic palate mesenchymal cells by down-regulation of TGF-beta/Smad signaling. Biochem Biophys Res Commun 340, 929–934, doi:10.1016/j.bbrc.2005.12.100 (2006).

77 Zhang, H. et al. Negative functional interaction of retinoic acid and TGF-beta signaling mediated by TG-interacting factor during chondrogenesis. Cell Physiol Biochem 23, 157–164, doi:10.1159/000204104 (2009).

78 Hu, Q. X. et al. All-trans-retinoic acid activates SDF-1/CXCR4/ROCK2 signaling pathway to inhibit chondrogenesis. Am J Transl Res 9, 2296–2305 (2017).

79 Wang, Y. G. et al. All-trans-retinoid acid (ATRA) suppresses chondrogenesis of rat primary hind limb bud mesenchymal cells by downregulating p63 and cartilage-specific molecules. Environ Toxicol Pharmacol 38, 460–468, doi:10.1016/j.etap.2014.07.008 (2014).

80 Zhang, T. G. et al. All-trans-retinoic acid inhibits chondrogenesis of rat embryo hindlimb bud mesenchymal cells by downregulating p53 expression. Mol Med Rep 12, 210–218, doi:10.3892/mmr.2015.3423 (2015).

81 Pan, H., Cai, N., Li, M., Liu, G. H. & Izpisua Belmonte, J. C. Autophagic control of cell ‘stemness’. EMBO molecular medicine 5, 327–331, doi:10.1002/emmm.201201999 (2013).

82 Taylor, R. C., Cullen, S. P. & Martin, S. J. Apoptosis: controlled demolition at the cellular level. Nature reviews. Molecular cell biology 9, 231–241, doi:10.1038/nrm2312 (2008).

83 Rodon, J., Dienstmann, R., Serra, V. & Tabernero, J. Development of PI3K inhibitors: lessons learned from early clinical trials. Nature reviews. Clinical oncology 10, 143–153, doi:10.1038/nrclinonc.2013.10 (2013).

84 Worby, C. A. & Dixon, J. E. Pten. Annual review of biochemistry 83, 641–669, doi:10.1146/annurev-biochem-082411-113907 (2014).

85 Masia, S., Alvarez, S., de Lera, A. R. & Barettino, D. Rapid, nongenomic actions of retinoic acid on phosphatidylinositol-3-kinase signaling pathway mediated by the retinoic acid receptor. Mol Endocrinol 21, 2391–2402, doi:10.1210/me.2007-0062 (2007).

86 Busada, J. T. et al. Retinoic acid regulates Kit translation during spermatogonial differentiation in the mouse. Dev Biol 397, 140–149, doi:10.1016/j.ydbio.2014.10.020 (2015).

87 Lopez-Carballo, G., Moreno, L., Masia, S., Perez, P. & Barettino, D. Activation of the phosphatidylinositol 3-kinase/Akt signaling pathway by retinoic acid is required for neural differentiation of SH-SY5Y human neuroblastoma cells. The Journal of biological chemistry 277, 25297–25304, doi:10.1074/jbc.M201869200 (2002).

88 Piskunov, A. & Rochette-Egly, C. A retinoic acid receptor RARalpha pool present in membrane lipid rafts forms complexes with G protein alphaQ to activate p38MAPK. Oncogene 31, 3333–3345, doi:10.1038/onc.2011.499 (2012).

89 Schoen, C. et al. MicroRNAs in Palatogenesis and Cleft Palate. Frontiers in physiology 8, 165, doi:10.3389/fphys.2017.00165 (2017).

90 Li, J. et al. Assessment of differentially expressed plasma microRNAs in nonsyndromic cleft palate and nonsyndromic cleft lip with cleft palate. Oncotarget 7, 86266–86279, doi:10.18632/oncotarget.13379 (2016).

91 Grassia, V. et al. Salivary microRNAs as new molecular markers in cleft lip and palate: a new frontier in molecular medicine. Oncotarget 9, 18929–18938, doi:10.18632/oncotarget.24838 (2018).

92 Walker, S. E., Spencer, G. E., Necakov, A. & Carlone, R. L. Identification and Characterization of microRNAs during Retinoic Acid-Induced Regeneration of a Molluscan Central Nervous System. Int J Mol Sci 19, doi:10.3390/ijms19092741 (2018).

93 Nervi, C. & Grignani, F. RARs and microRNAs. Subcell Biochem 70, 151–179, doi:10.1007/978-94-017-9050-5_8 (2014).

94 Lima, L. et al. Modulation of all-trans retinoic acid-induced MiRNA expression in neoplastic cell lines: a systematic review. BMC Cancer 19, 866, doi:10.1186/s12885-019-6081-7 (2019).

95 Garzon, R. et al. MicroRNA gene expression during retinoic acid-induced differentiation of human acute promyelocytic leukemia. Oncogene 26, 4148–4157, doi:10.1038/sj.onc.1210186 (2007).

96 Liu, B. et al. Retinoid acid-induced microRNA-31-5p suppresses myogenic proliferation and differentiation by targeting CamkIIdelta. Skelet Muscle 7, 8, doi:10.1186/s13395-017-0126-x (2017).

97 Li, Z. et al. A microRNA signature for a BMP2-induced osteoblast lineage commitment program. Proceedings of the National Academy of Sciences of the United States of America 105, 13906–13911, doi:10.1073/pnas.0804438105 (2008).

98 Gao, J. et al. MicroRNA expression during osteogenic differentiation of human multipotent mesenchymal stromal cells from bone marrow. Journal of cellular biochemistry 112, 1844–1856, doi:10.1002/jcb.23106 (2011).

99 Li, H. et al. miR-17-5p and miR-106a are involved in the balance between osteogenic and adipogenic differentiation of adipose-derived mesenchymal stem cells. Stem Cell Res 10, 313–324, doi:10.1016/j.scr.2012.11.007 (2013).

100 Jia, B. et al. A feed-forward regulatory network lncPCAT1/miR-106a-5p/E2F5 regulates the osteogenic differentiation of periodontal ligament stem cells. Journal of cellular physiology 234, 19523–19538, doi:10.1002/jcp.28550 (2019).

101 Manochantr, S. et al. The Effects of BMP-2, miR-31, miR-106a, and miR-148a on Osteogenic Differentiation of MSCs Derived from Amnion in Comparison with MSCs Derived from the Bone Marrow. Stem Cells Int 2017, 7257628, doi:10.1155/2017/7257628 (2017).

